# Immature neurons in the primate amygdala: changes with early development and disrupted early environment

**DOI:** 10.1101/2023.02.10.528076

**Authors:** Alexandra C McHale-Matthews, Danielle M DeCampo, Tanzy Love, Judy L Cameron, Julie L Fudge

**Author notes:** Correspondence to: Julie L. Fudge, M.D., Departments of Neuroscience and Psychiatry The Del Monte Institute for Neuroscience University of Rochester Medical Center. **Email addresses**: Alexandra McHale-Matthews; Danielle DeCampo; Tanzy Love; Judy Cameron; Julie Fudge.

## Abstract

In human and nonhuman primates, the amygdala paralaminar nucleus (PL) contains immature neurons. To explore the PL’s potential for cellular growth during development, we compared PL cells in 1) infant and adolescent macaques (control, maternally-reared), and in 2) infant macaques that experienced separation from their mother in the first month of life. In maternally-reared animals, the adolescent PL had fewer immature neurons, more mature neurons, and larger immature soma volumes compared to infant PL. There were also fewer total neurons (immature plus mature) in adolescent versus infant PL, suggesting that some neurons move out of the PL by adolescence. Maternal separation did not change mean immature or mature neuron counts in infant PL. However, across all infant animals, immature neuron soma volume was strongly correlated with mature neuron counts. *tbr-1* mRNA, a transcript required for glutamatergic neuron maturation, is significantly reduced in the maternally-separated infant PL (DeCampo et al, 2017), and was also positively correlated with mature neuron counts in infant PL. We conclude that immature neurons gradually mature by adolescence, and that the stress of maternal separation may shift this trajectory, as revealed by correlations between tbr1mRNA and mature neuron numbers across animals.

## Introduction

The amygdala is a heterogeneous collection of nuclei important for social-emotional processing, and is dysregulated in various neuropsychiatric diseases (Klüver & Bucy, 1939; LeDoux, 2000; Nishijo, Ono, & Nishino, 1988; Schumann, Bauman, & Amaral, 2011). One little-known amygdala nucleus, which is prominent in long-lived mammalian species, is the paralaminar nucleus (PL). The PL is a cellular region that surrounds the basal nucleus in postnatal monkeys and humans, reaches its largest size in humans, and persists over development (Review, (D. de Campo & Fudge, 2011)). Originally thought to be a small-celled portion of the basal nucleus (Braak & Braak, 1983; Jimenez-Castellanos, 1949), and largely ignored, the PL is now known to be a developmentally distinct region from the basal nucleus and contains immature neurons that persist throughout life (Avino et al., 2018; Chareyron, Amaral, & Lavenex, 2016; D. M. de Campo et al., 2017; Fudge, 2004; Sorrells et al., 2019). Immature neurons in both the adult monkey and human PL stain abundantly for markers important for post-mitotic neural differentiation and migration such as doublecortin (DCX), B-cell lymphoma (Bcl-2), polysialated neural cell adhesion molecule (PSA-NCAM), and beta-tubulin III (TUJI). Although originally thought to emerge from the former ganglionic eminence (Ulfig, Setzer, & Bohl, 2003), the human PL is now known to originate from the lateral or ventral pallium since its immature neurons express pallial protein markers associated with a glutamatergic fate (*tbr1* and *VGluT 2*, among others) (Sorrells et al., 2019).

The presence of immature-appearing neuroblasts in the PL implies a substrate for increases in mature neurons over postnatal life. Chareyron and colleagues provided the first evidence that the PL may contain a pool of immature neurons that are poised for differentiation and migration over a protracted developmental window (Chareyron, Lavenex, Amaral, & Lavenex, 2012). They counted immature-appearing and mature neurons using morphologic criteria (cresyl violet staining) in the PL of infant monkeys and older animals, and reported that immature neuron counts declined, mature neuron counts increased, and that average soma volume of PL neurons increased with age. However, a limitation of this study was assessment of putative immature and mature neuronal changes using cellular morphology alone. Furthermore, while developmental timepoints through the first year of life were thoroughly investigated in that study, pooled data from 5-9 year old monkeys was used, due to small samples of older animals (Chareyron et al., 2012). To bridge these gaps, and focus on adolescent development, we assessed immature neurons using DCX immunoreactivity (DCX-IR) to examine whether there were signs of neuronal differentiation in the PL of female primates between infant (3-months of age, equivalent to 9-12 month old humans) and adolescent (4-years of age, equivalent to 16 year old humans) time points. These time points were selected because of potentially dramatic developmental shifts between these age groups, based on human studies (Avino et al., 2018; Sorrells et al., 2019). In the first part of this study, we specifically investigated how the number of immature (DCX-positive) neurons, the number of mature (DCX-negative) neurons, and the soma volume of immature (DCX-positive) neurons changed with age to understand what broad, cellular alterations might occur between these two time points.

Postnatal maturation of neuronal precursor growth in the PL can potentially be affected by early life environmental perturbations (D. M. de Campo et al., 2017). Early life maternal separation is a well-studied environmental perturbation since interactions with primary caregivers, or lack thereof, strongly influences long-term behavior and health across many mammalian species (Harlow, Dodsworth, & Harlow, 1965; Sabatini et al., 2007; Sullivan et al., 2006; Suomi, Collins, & Harlow, 1973). Examples of behavioral changes that follow maternal separation in monkeys and humans include indiscriminate social approach, less play and locomotor activity, and fewer distress calls than before deprivation occurs (Hinde, Spencer-Booth, & Bruce, 1966; O’Connor & Rutter, 2000). Strikingly, these behavior changes are specifically correlated with gene expression (monkey) and connectional (human) abnormalities in the amygdala (D. M. de Campo et al., 2017; Gee et al., 2013; Sabatini et al., 2007; Tottenham et al., 2010). We sought to address translational questions between these species by focusing on cellular changes associated with maternal separation in female infant monkeys.

Since traditional maternal separation paradigms may not fully mimic typical human maternal separation (Harlow et al., 1965; Suomi et al., 1973), we used a more ethological model (J.L. Cameron, 2001; McCormick, Gualano, Kerr, Rockcastle, & Cameron, 2005; Sabatini et al., 2007). In this model, the mother is removed from the social group of juvenile and adolescent animals, leaving other older juveniles and adolescents to care for the infant (as occurs in the wild) (J. L. Cameron, Eagleson, Fox, Hensch, & Levitt, 2017; D. M. de Campo et al., 2017; Howell et al., 2017; Sabatini et al., 2007). Previously examining the same cohort of monkeys used in this study, we found that by 3 months of age, one of the most downregulated gene transcripts in the PL of separated infants was *tbr-1* (D. M. de Campo et al., 2017), a critical transcription factor for the maturation of glutamatergic precursors (Bulfone et al., 1995; S. F. Darbandi et al., 2020; Hevner et al., 2001). However, the cellular correlates of decreased *tbr-1* transcript levels in separated infants were unclear. Therefore, as part of the present study, we used immersion fixed tissue from the opposite hemisphere in this same cohort to examine potential cellular alterations in maternally-separated infants by 3 months of age.

## Materials & Methods

### Animals

Animals used were from the same cohorts born and reared at the University of Pittsburgh, and used in previous studies (D. M. de Campo et al., 2017; Sabatini et al., 2007). We approached analyses into two parts: 1) cellular comparisons between control infant and adolescent female macaque cohorts (Normal development), and 2) analyses examining cellular comparisons between maternally-reared female infant macaques, and 2 groups of maternally-separated female infants (Maternal separation in infants) (Fig. 1). All experiments were approved by the University of Pittsburgh Animal Care and Use committee and were in accordance with guidelines issued by the National Institute of Health.

**Figure 1:**
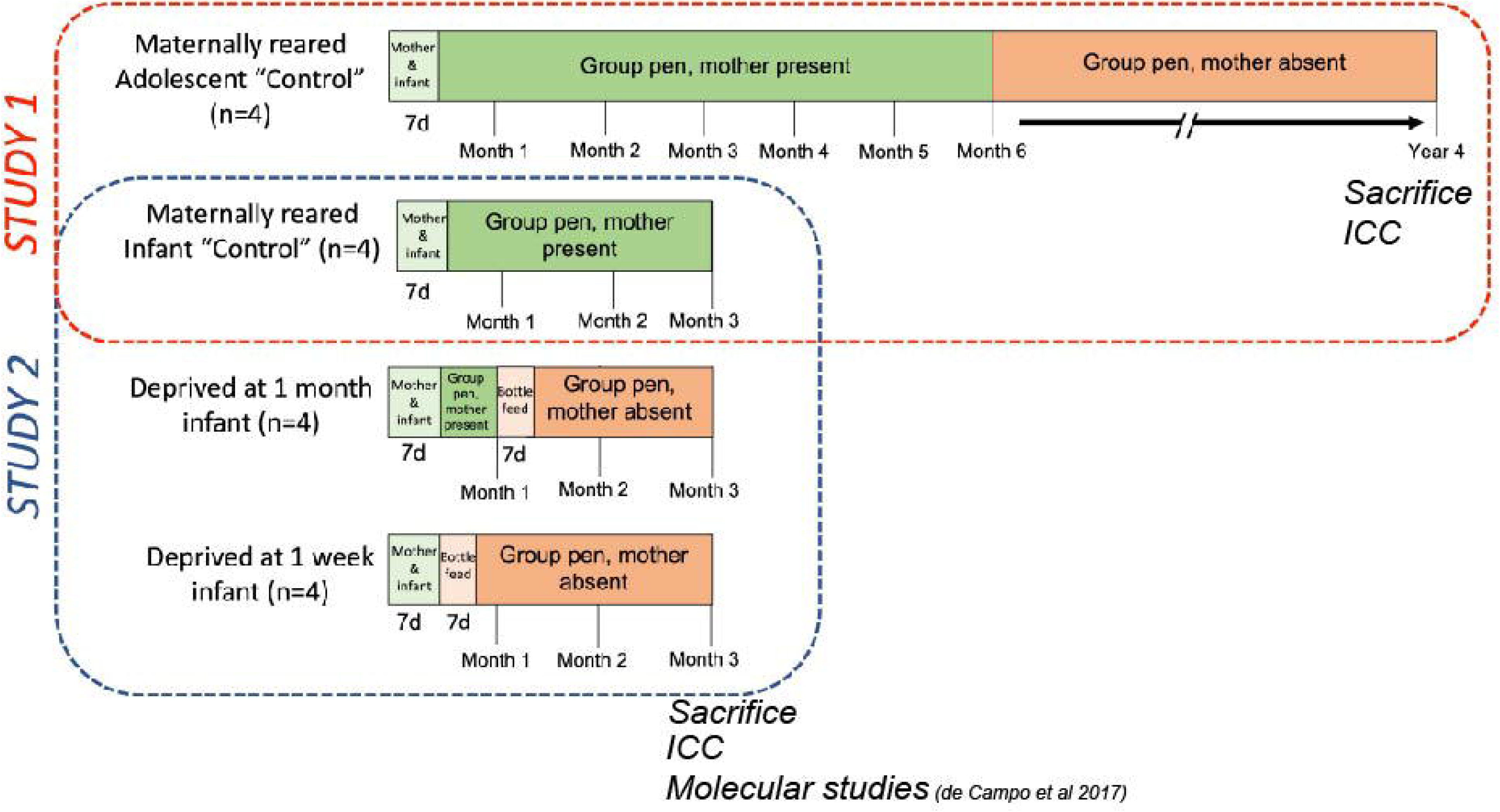
Monkey housing paradigms for Normal Postnatal Development (Study 1) and Maternal Separation (Study 2). “Study 1” (outlined in red) examined a normal development paradigm and consisted of 2 groups: a “maternally reared adolescent control” group (top panel), and a “maternally reared infant control” group with n = 4 animals per group. Panels are organized from left to right chronologically from birth to time of sacrifice (4 years of age for adolescent control; 12 weeks of age for infant control). Green bars indicate that the mother is present during that time period (light green indicates that the infant is singly housed with the mother, and dark green indicates that the infant is in a group rearing paradigm with the mother present). The dark orange bar indicates that the mother is not present during that time period (in adolescent control, this is the typical age at which the mother leaves the infant to form a consort with a male macaque in the next mating season). “Study 2” (outlined in blue) examined an early-life maternal separation paradigm and consisted of 3 groups: a “maternally-reared control” group (also analyzed in “Study 1”), a “separated at 1-month” group, and a “separated at 1-week” group, n = 4 animals per group. Orange bars indicate that the mother is not present during that time period (light orange indicates the 7-day period in which the infant is singly housed in a cage adjacent to the group-rearing pen to learn how to bottle feed).

### Rearing environment

All animals shared a single cage with their mother during the first week of life (per typical animal husbandry protocols) and were then randomly assigned to an experimental group (Fig. 1). Control animals were introduced to a group-rearing pen environment with their mother present for the duration of the study for infants (week 2 through week 12), and through 6 months of age for maternally-reared adolescents. For this latter group, the mother was removed when the adolescent animals were 6 months of age, since this is the typical point at which macaque mothers leave their infant and form a consort with a male macaque for the next mating season (J. L. Cameron et al., 2017). Adolescent monkeys then remained in the same group pen without their mother until 4 years of age.

Group-rearing pen environments contained 4-5 primates ranging in age, including an adolescent female due to their particular attentiveness to infants. Only 1 experimental infant and its mother were present in each group-rearing pen at a time. When present, the mother was always the most dominant member of the group-rearing pen to insure there was no added stress on the mother. Each group-rearing pen was an all-cement block construction measuring 14ft X 11ft X 12ft, and contained a chain-link pen-front, four perches at varying heights, and a thick layer of wood shavings on the floor. Full details of the animal husbandry protocol are available in (Sabatini et al., 2007).

### Maternal separation paradigm (Fig. 1)

For the maternal separation portion of the study, brain tissue from three groups of infant animals sacrificed at 3 month of age (n = 4 animals per group; 12 animals total, all female) was assessed: a control (maternally-reared group), and two groups that had their mother removed from their group-rearing environment at either 1-week of age (“separated at 1-week”) or 1-month of age (“separated at 1-month”). Complete details of the paradigm have been previously published (D. M. de Campo et al., 2017; Sabatini et al., 2007).

Briefly, all 3 infant groups shared a single cage with their mother during the first week of life, then were randomly assigned to a “separated at 1-week” group, a “separated at 1-month” group, or a “maternally-reared” infant control group. After the first week singly housed with the mother, the separated at 1-week infants had the mother removed from the cage and the infant was left in the single cage to learn how to bottle feed for a 5-7 day period (*ad libitum* Similac with Iron baby formula; Abbott Laboratories, Columbus OH). A soft, cotton-stuffed toy was placed in the cage to provide contact comfort. After this period, the separated at 1-week animal was introduced to that same group-rearing pen environment for weeks 3-12 without their mother present. A hutch with a door small enough for the separated infant to transgress, but too small for other animals to enter, was placed in the pen and bottles were made available to the infant in this hutch. Separated at 1-month infants were incorporated into a group-rearing pen environment with their mother present for weeks 2-3. Then at the start of week 4, the separated at 1-month infants spent a 5-7 day period in an adjacent cage with a cotton snuggly toy to learn how to bottle feed. After this period, they were re-introduced to that same group-rearing environment without their mother present for weeks 6-12. Again, a hutch with a small door was available in the pen after the mother was removed and bottles were made available in the hutch. Maternally-reared infant control animals were introduced to a group-rearing pen environment with their mother present for the duration of the study (week 2 through week 12).

### Surgery and brain-cutting protocols

Brain tissue was collected at 12 weeks of age (n=12, infant groups), or at 4 years of age (n=4, adolescent group) at the University of Pittsburgh. Monkeys were initially anesthetized with ketamine HCl (10 mg/kg, i.m. injection), then deeply anesthetized with sodium pentobarbital (30 mg/kg, i.v.). A transcardial perfusion surgery followed; the monkey’s chest was opened, and a catheter was placed in the heart such that the catheter tip was located in the ascending aorta. Approximately 1 liter of ice cold 0.9% NaCl solution containing 2% sodium nitrite and 5000 IU heparin was transcardially perfused. Perfusate ran out of the opened left atrium, and the monkey was quickly decapitated once the perfusate was clear. The brain was then promptly removed from the skull and hemisected on the mid-sagittal plane. The left hemisphere was cut in coronal blocks and immersion fixed. The right hemisphere was flash frozen and used for gene expression studies as previously described (D. M. de Campo et al., 2017; Sabatini et al., 2007).

Fixed blocks were submerged into 4% paraformaldehyde (PFA), then placed in a 3:3:3:1 ratio of Glycerol, Ethylene glycol, distilled H_2_O, and 2X phosphate buffer (PB) for storage in a −20°C freezer. For this study, blocks were selected and cut approximately at the decussation of the anterior commissure anteriorly, and at the level of the midbrain caudally to capture the rostro-caudal extent of the amygdala and PL.

Cryoprotected blocks were transferred daily into fresh 18% sucrose solution (sucrose in 1:9:10 ratio of 10X saline, distilled and dionized H_2_O, and 2X PO_4_ buffer) over a 14-day period. Then, blocks that included the rostral-caudal extent of the amygdala were sectioned coronally on a freezing, sliding microtome at 40 μm. Brain slices were placed in 12 consecutive compartments containing cold cryoprotectant solution (30% sucrose and 30% ethylene glycol in PB), then stored at −20°C (Rosene, Roy, & Davis, 1986). 1:12 sections were the chosen sampling increment based on our condition-setting stereological studies in normal adolescent animals (not part of this study), and with reference to Chareyron et al (Chareyron et al., 2012). Compartments for each animal contained evenly spaced sections with a random starting point. All staining batches were counterbalanced in a blinded manner.

### Histology for fixed tissue

#### Doublecortin (DCX)

Conditions for DCX immunostaining were first established in control animals that were not part of this study (Fudge, deCampo, & Becoats, 2012). Tissue was rinsed in PB with 0.3% Triton-X (PB-TX) overnight. The next day, brain slices were treated with an endogenous peroxidase inhibitor for 5 minutes, and then underwent more rinses in PB-TX. Sections were then pre-incubated for 30 minutes in 10% normal goat serum blocking solution with PB-TX (NGS-PB-TX). All sections were then incubated in primary antisera to DCX (1:15000, Abcam, rabbit), at 4°C for four nights. Sections were then thoroughly rinsed, blocked with 10% NGS-PB-TX, and incubated for 40 minutes in the appropriate biotinylated secondary antibody. After more rinses, sections with bound anti-DCX antibodies were incubated in an avidin-biotin complex (Vectastain ABC Elite; Vector Laboratories), visualized with 3,3’-Diaminobenzidine (DAB), and then activated with 0.3% hydrogen peroxide (H_2_O_2_).

#### Cresyl Violet (Nissl)

DCX-stained sections were mounted onto gelatin coated slides from 0.1M PB solution and air-dried over a 2- to 4-week period. They were then rapidly dehydrated and rehydrated, and counterstained with a light cresyl violet stain (Chroma-Gesellschaft; West Germany), and coverslipped with DPX Mounting Medium (Electron Microscopy Sciences; Hatfield, PA). In order to maintain the section height required for optical fractionator analyses, slight modifications to the Nissl-staining protocol were made to minimize dehydration from ethanol.

#### Acetylcholinesterase (AChE)

AChE staining in adjacent sections was also used to distinguish between basal nucleus subdivisions and the PL (Amaral & Bassett, 1989), using the Geneser-Jensen technique (Geneser-Jensen & Blackstad, 1971).

### Stereological methods

All analyses were conducted in a blinded manner, where all identifying information on each slide was masked, and a number was randomly assigned to each monkey to shield its group identity. 1:12 sections double-labeled for DCX and Nissl were examined with an Olympus UPlanFL 100x/1.30 oil lens using an Olympus AX70 microscope interfaced with Stereoinvestigator via a video CCD (Microbrightfield, Williston, VT). On average, 8 sections per animal were examined (range = 7-9 sections) with approximately 480μm of distance between each slide. Our analysis included the PL where it first appeared ventral to the basal nucleus. The rostral-most portion of the PL surrounds the anterior basal nucleus, forming a DCX-positive neuronal “cap”, but is not present in every series. In order to prevent an over-representation of cells in some monkeys compared to others, we excluded this anterior-most portion of the PL from analysis.

#### Optical fractionator

Neuron counts were estimated using a well-established, unbiased, and systematic random sampling method known as the optical fractionator (H. J. G. Gundersen & Jensen, 1987; West, Slomianka, & Gundersen, 1991) (Stereoinvestigator, Microbrightfield Biosciences, Williston, VT). Briefly, the PL was outlined under low power magnification (2x objective) (cyan line, Fig. 2A). The boundaries of the PL were determined by a clear density of DCX-positive neurons seen at 2x magnification in the region ventral to the basal nucleus. In the software, a sampling grid with evenly spaced counting frames is randomly placed on the region of interest (Fig. 2A & 2B). Sampling in each counting frame occurs under high power (100x oil objective) (Fig. 2C, Table 1). Counting frames, and labeled cells associated with them, occasionally fall upon the boundaries of the contour for some sections, according to the random placement of the grid across many sections over the course of a study (Fig. 2C). Rules for counting labeled neurons apply to all counting frames regardless of whether they fall on contour boundaries. All DCX-positive neurons were counted as “immature” neurons, regardless of morphology; “mature” neurons were defined as DCX-negative cresyl violet stained cells with characteristic nuclear staining (Chareyron et al., 2012; Garcia-Cabezas, John, Barbas, & Zikopoulos, 2016). Sampling parameters (counting frame and scanning grid dimensions) were first established in a preliminary study neural population by oversampling to yield a coefficient of error (CE) of 0.10 or lower in each population for both ages (Table 1). Then, cases were re-coded and blindly counted. Section thickness was collected at every sampling site, so that the final cell estimates were calculated using “number weighted section thickness”.

**Figure 2:**
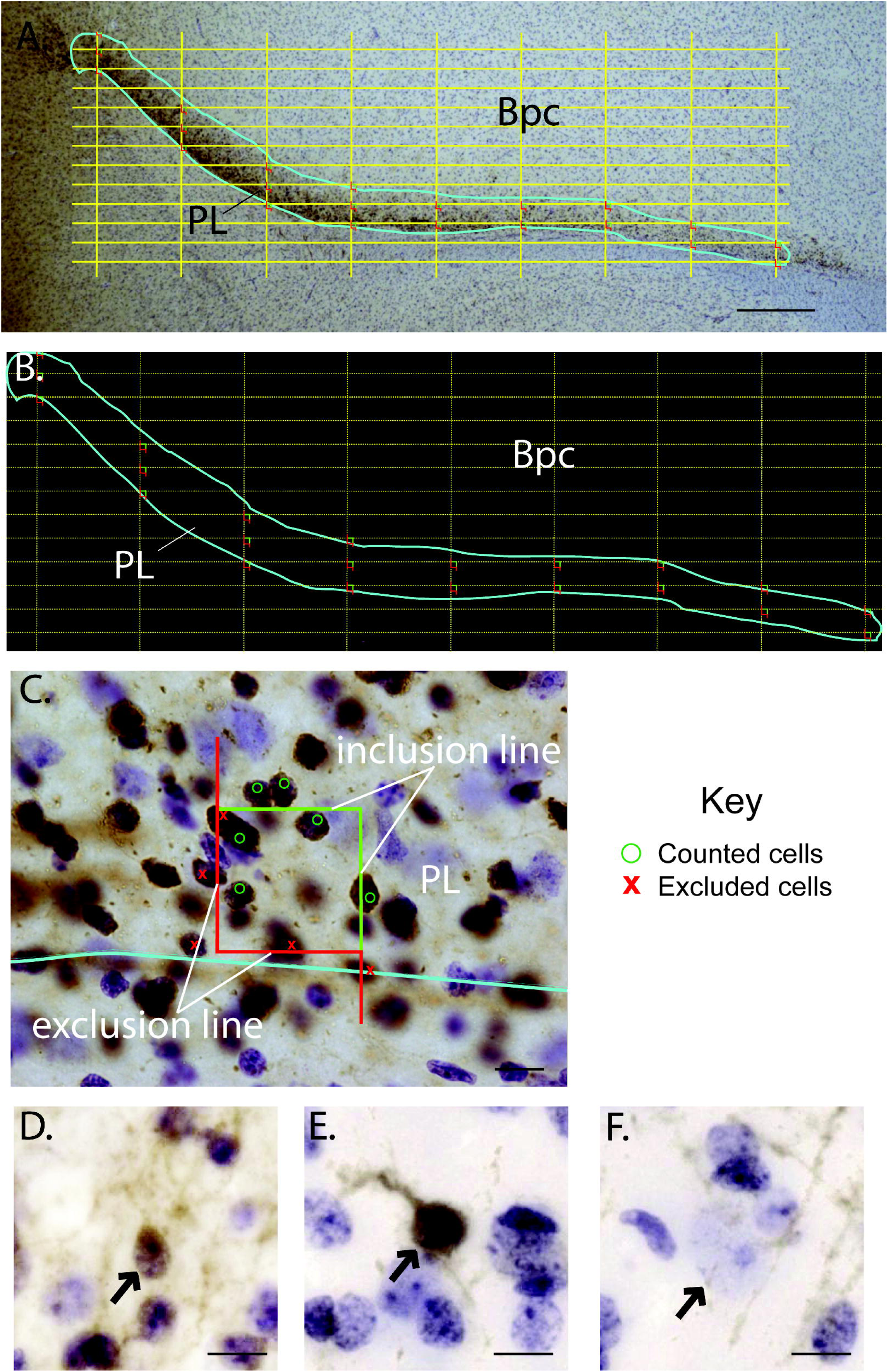
Stereology study design. **A.-B.** Coronal section of the ventral amygdala at 2 magnifications, with paralaminar nucleus (PL) outlined in cyan, and grid dimensions (yellow dotted lines, 545 μm x 124 μm) with counting frame (boxes containing green and red lines) overlaid. All counting frames at least partially touch the region of interest, and all data within these counting frames are considered valid for counting purposes. **C.** Higher power view of a counting frame. The green lines are “inclusion lines”, and red lines in the counting frame indicates an “exclusion line” (West, Slomianka, & Gundersen, 1991). 2x magnification, scale bar = 500 μm. Per sampling criteria, DCX-positive cells marked with green “O”, and DCX-positive cells marked with red ‘X’ are excluded (see Methods) 100x magnification, scale bar = 10 μm. **D-F.** Brightfield images of “small” DCX-positive (immature) neuron (**D.**), “large” DCX-positive (immature) neuron (**E.**), and DCX-negative (mature) neuron nucleus stained with cresyl violet (**F.**), 40x magnification, scale bar = 10 μm. Bpc, parvicellular basal nucleus; PL, paralaminar nucleus.

**Table 1:** Stereological parameters used for infant and adult tissue.

| **Age and cell groups** | **Counting frame (μm)** | **Scan grid (μm)** | **Disector height (μm)** | **Guard zones (μm)** | **Average measured section thickness (μm)<sup>2</sup> (including range)** | **Average number of cells counted (including range)** | **Average C.E. (including range)** |
| --- | --- | --- | --- | --- | --- | --- | --- |
| Infant DCX+ neurons | 30 x 30 | 545 x 124 | 10 | 2 | 17.78 (12.6 – 26.6) | 586 (381 - 810) | 0.05 (0.04 – 0.06) |
| Infant Mature Nissl-stained neurons | 60 x 60 | 545 x 124 | 10 | 2 | 17.60 (12.8 – 24.7) | 225 (100 – 401) | 0.07 (0.05 – 0.10) |
| Adolescent DCX+ neurons | 30 x 30 | 545 x 124 | 10 | 2 | 18.78 (14.3 – 22.8) | 349 (281 – 408) | 0.06 (0.05 – 0.06) |
| Adolescent Mature Nissl-stained neurons | 60 x 60 | 545 x 124 | 10 | 2 | 18.42 (13.8 – 22.5) | 535 (420 – 642) | 0.05 (0.04 – 0.05) |

#### Cell counting criteria

As per standard criteria (H. J. G. Gundersen & Jensen, 1987; West et al., 1991), all counting frames systematically sample the region of interest, including “border” zones (called the “contours”, or the perimeter). Z-height was measured for each counting frame. For DCX-positive cells, we established that the cell soma (i.e. nucleus and surrounding cytoplasmic rim) had to be either within the counting frame, or if touching the green “inclusion boundary”, at least partially inside the disector. Mature neurons were identified by lack of DCX-IR as well as characteristic Nissl-stained nuclei: the characteristic large pale nucleus and small, dark nucleolus found in mature pyramidal cells. The DCX-negative nucleus (“mature”) had to be fully within the disector boundaries, and if touching the green “inclusion” line, at least partially inside the disector. Regardless, the nucleolus needed to be in focus within the disector Z-plane to be counted (Fig. 2C). If the cell body (or nucleus, in the case of DCX-negative mature neurons) was wholly outside of the disector, or was inside but touched/crossed the exclusion line (red) of the disector, it was not counted. These criteria prevent double counting of cells in the next sampling box on closely spaced grids (Slomianka & West, 2005).

#### Nucleator method

While collecting immature cell counts, we simultaneously implemented the nucleator method, a method for calculating soma volume, on immature DCX-positive neurons at each counting frame (H. J. Gundersen, 1988). For the nucleator method, we placed a series of 4 rays that emanated from the nucleolus and marked where these rays intersected with the outer boundary of the soma to capture the average radius of that cell. Then, the formula 4/3π(mean length of rays)^3^ was used to calculate the DCX-positive soma volume, assuming sphericity of the cell. The soma volume of every DCX-positive neuron that met optical fractionator sampling criteria was measured.

#### In situ hybridization for tbr-1 (conducted in a previous study (DeCampo et al, 2017))

For assessing *tbr-1* mRNA expression in the maternal separation study, we used the right hemisphere (flash frozen) from the infant animals, which had been stored at −80°C (D. M. de Campo et al., 2017; Sabatini et al., 2007). Using a cryostat, 20μm-thin slices were sectioned through the caudal amygdala containing the PL. Ten adjacent sections of the caudal amygdala (containing the PL) per animal were placed onto emulsion-dipped slides, then processed for in situ hybridization of *tbr-1* mRNA. Photomicrographs for these 10 sections per animal were collected and analyzed as 8-bit grayscale images using Image J64. To count silver grains, a pre-specified frame measuring 646 μm x 136.24 μm was placed over the PL. The same frame was placed on an adjacent non-tissue area to determine background levels. Silver grains were identified using the “IJ isodata” function, then counted if they had an area ranging from 2.58 μm^2^ – 5.37 μm^2^ when using the “Analyze Particles” function. The four slides with the highest number of silver grains were identified, then the mean number of silver grains from these four slides per animal were calculated, with background subtracted.

### Statistical analyses

#### Normal development study (Study 1)

Shapiro-Wilk test for normality and Bartlett tests for homogeneity of variance were conducted for DCX-positive neuron count estimates, DCX-negative neuron count estimates, summed DCX-positive neuron count estimates, DCX-positive dorsal/ventral cell count estimates, DCX-negative neuron count estimates, ratio of DCX-positive neuron count estimates to DCX-negative neuron count estimates, and DCX-positive neuron soma volume, and were found to be normally distributed with equal variance across groups, respectively, except soma volumes. One-tailed, two-sample t-tests were then used to examine differences in a specific direction, along development trajectories (e.g. greater soma size in older animals). DCX-positive soma volume data (0.5% trimmed means to eliminate improbably small and large cells) was set to display the range of DCX-positive soma volumes in infant monkeys and adolescent monkeys (RStudio packages “ggplot2”, “ggpubr”, “forcats”, and “ggprism”, and binwidth set to 10). We used the median DCX-positive soma volume in the control infant animal group as the break-point between “small” or “large” neuron DCX-positive soma volumes. Since Shapiro-Wilks test revealed non-normally distributed data, one-tailed, unpaired, two-samples Wilcoxon Rank Sum tests were run for DCX-positive soma volumes, “small” DCX-positive soma volumes, and “large” DCX-positive soma volumes separately. If p<0.05, findings were considered significant.

#### Maternal separation in infants study (Study 2)

Shapiro-Wilk and Bartlett’s tests for DCX-positive neuron count estimates, DCX-positive dorsal/ventral cell count estimates, DCX-negative (mature) Nissl-labeled neuron count estimates, summed DCX-positive neuron count estimates and DCX-negative neuron count estimates, and ratio of DCX-positive neuron count estimates and DCX-negative neuron count estimates revealed these data were normally distributed with homogeneity of variance, respectively, and one-way ANOVA tests were run (alpha set at 0.05). Tukey Honestly Significant Differences (Tukey HSD) with Benjamini-Hochberg (BH) correction post-hoc tests were conducted to perform pairwise comparisons between group means. Pairwise comparisons were considered significant if the p-value associated with it was < 0.05. Bartlett’s p-value for average DCX-positive neuron soma volume was less than 0.05 and the data did not have homogeneity of variance. Therefore, we applied non-parametric Kruskal-Wallis one-way ANOVA. The tests were considered significant if the p-value was < 0.05, and then Dunn’s test were conducted for post hoc comparisons with Benjamini–Hochberg (BH) correction.

DCX-positive soma volume data was analyzed similar to that in Study 1 (above) for three infant groups, again using 0.5% trimmed means. DCX-positive soma volume was divided at the “infant control” median break-point, to classify “small” neurons, and “large” neurons falling above and below the median, respectively. Shapiro-Wilks test again revealed lack of normality for these data. Kruskal-Wallis one-way ANOVA tests were performed for DCX-positive soma volume, “small” DCX-positive soma volume, and “large” DCX-positive soma volume separately, for each outcome. Dunn’s test was conducted for post hoc comparisons with Benjamini–Hochberg (BH) correction.

#### Relationships between immature and mature neuron cell counts, immature neuron soma size, and tbr-1 mRNA levels

Pearson’s correlations were used to examine relationships between average immature DCX-positive soma volumes and 1) DCX-positive immature neuron count and 2) DCX-negative mature neuron count. Pearson’s correlations were also used to examine relationships between *tbr-1* silver grain counts (previous dataset, (D. M. de Campo et al., 2017)) versus 1) immature DCX-positive neuron count, 2) mature DCX-negative neuron count, and 3) average immature DCX-positive neuron soma volume. All analyses were conducted in RStudio using the “ggpubr” package. One-tailed *p* values with n-2 degrees of freedom were used. Correlations were considered significant if the p-value associated with it was < 0.05. All statistical analyses including Rmarkdown code are available at https://figshare.com/s/370067d5836d5.

## Results

### Overall findings

#### Morphological and cytological features of cell types in the PL (Figs. 2D-F)

Immature neurons ranged from small, round profiles (Fig. 2D) to a larger, more differentiated appearance with thicker processes (Fig. 2E). Immature neuron somata were ovoid, circular, or sometimes tear-drop shaped, with a nucleolus and 1-2 heterochromatin bundles. All DCX-negative cell types were identified by their morphology using cresyl violet staining, and were classified based on previously established criteria (Chareyron et al., 2012; Garcia-Cabezas et al., 2016). Mature (DCX-negative) neurons in Nissl stain were the largest cells in the region, and had very lightly stained nuclei, with occasional specks of chromatin, and a prominent, medium/large-sized nucleolus (Fig. 2F). For reference, astrocytes were smaller than mature neurons and had generally “light” nuclear staining with several scattered bundles of heterochromatin appearing in the cytoplasm. Oligodendrocytes were smaller than astrocytes, round, had dark nuclear staining, and tended to appear in bundles. Microglia stained darkly, had a heterogeneous cell shape, and could be comma-shaped, rod-shaped, round, or ameboid (not shown). These cell types were not counted.

#### Heterogeneity of immature neurons in the PL

The PL was characterized by very packed, small DCX-positive cells in both adolescent and infant groups (Fig. 3A-D, region between cyan lines). Traveling dorsally toward the Bpc in all animal groups, a greater heterogeneity in DCX-positive neuron size and morphology was observed in the dorsal third of the structure (yellow dotted line), including scattered pyramidal-shaped DCX-positive cells with thick proximal dendrites along with smaller, round DCX-positive soma (Fig. 3, white stars). Even within “small” and “large” categories of DCX-positive neurons, a range of soma sizes was appreciated in analyses using 100x oil magnification.

**Figure 3:**
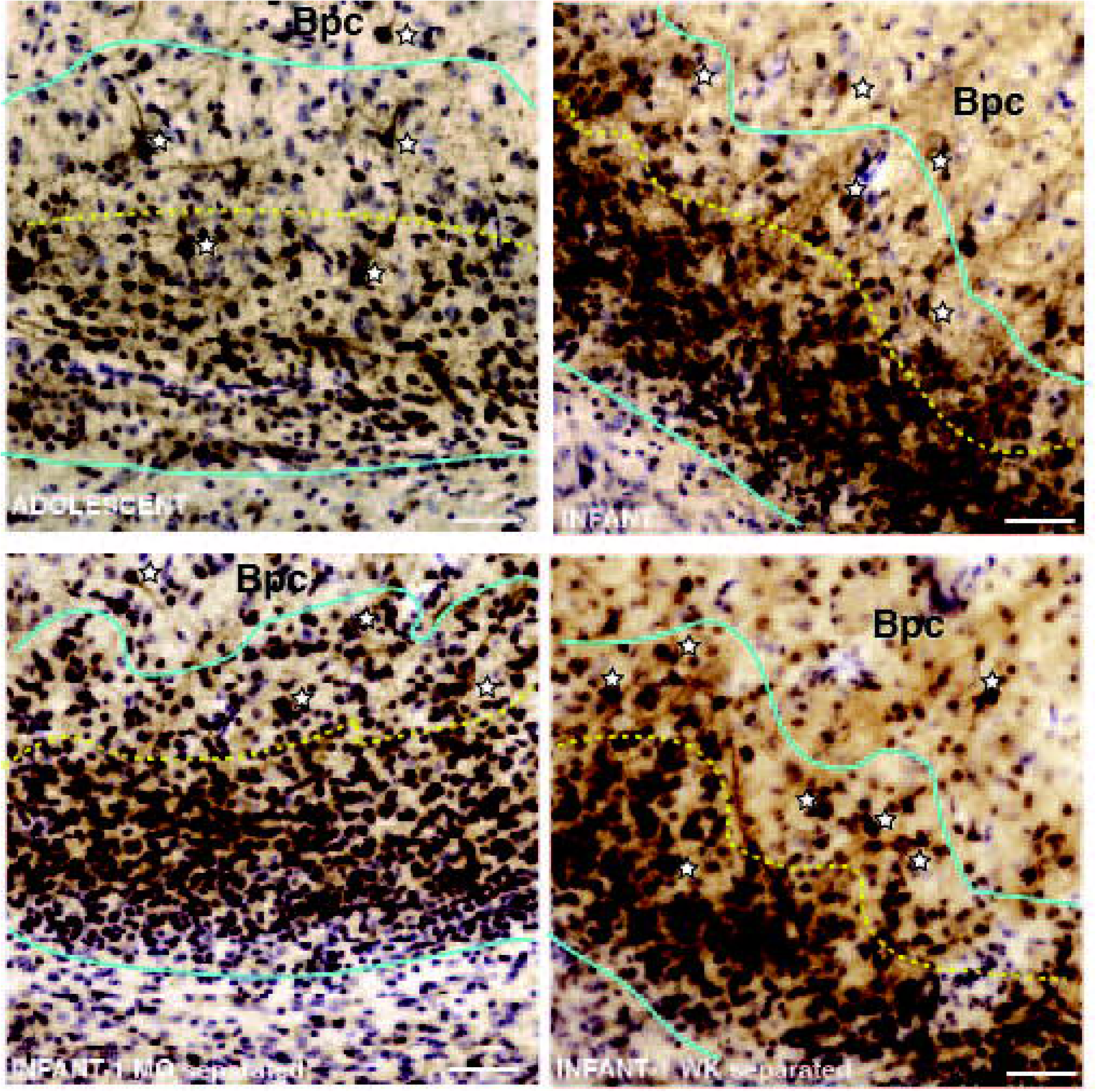
DCX-positive neuron size differences in the ventral-dorsal extent of the PL. **A.** Adolescent PL, outlined in cyan under low power, showing morphologic heterogeneity of DCX neurons, with many small round DCX-positive cells in the ventral PL, and more heterogeneous morphologies, including larger DCX-positive neurons with proximal dendrites, in the dorsal third (above the yellow dotted line, white stars). Some larger DCX-positive neurons were also scattered in the overlying Bpc. Similar images for: **B.** Infant maternally reared control, **C.** Separated at 1 month infant, and **D.** Separated at 1 week infant. Scale bar = 50 μm. Bpc, parvicellular basal nucleus; PL, paralaminar nucleus.

### Normal Postnatal Development from infancy to adolescence (STUDY 1)

#### Cellular measures in infant and adolescent macaques

##### Immature (DCX-positive) neuron count estimates (Fig. 4A)

There was an average of 907,008 DCX-positive (sample standard deviation (SD) = 188,260.05) neurons present in infant monkeys, ranging from 678,985 neurons - 1,129,609 neurons. There was an average of 541,165 DCX-positive (SD=95,070.68) neurons present in adolescent monkeys, ranging from 446,114 neurons - 668,092 neurons. Differences between these groups were statistically significant (p=0.007), with the number of immature neurons in the adolescent PL comprising 60% of that in the infant.

**Figure 4:**
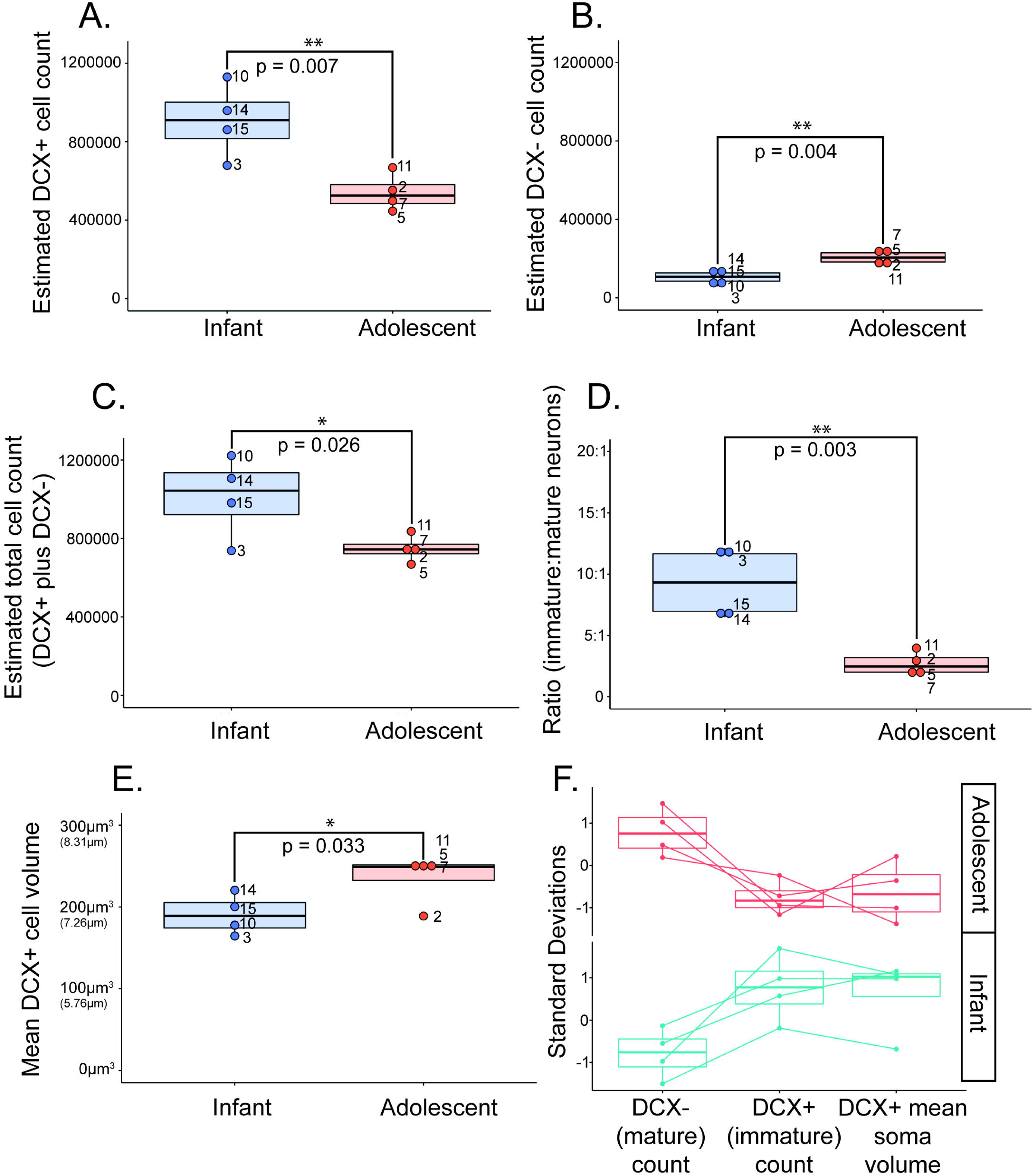
Normal Postnatal Development study: Cellular measurements. Boxplots showing infant (blue) and adolescent (orange) data for **A.** DCX-positive (immature) neuron count estimates, **B.** DCX-negative (mature) neuron count estimates, **C.** Summed DCX-positive (immature) and DCX-negative (mature) neuron counts, **D.** Ratio of DCX-positive (immature) to DCX-negative (mature) neuron counts, and **E.** average DCX-positive neuron soma volume data (μm^3^) and its corresponding diameter (μm). Numbers next to data points represent for individual monkeys in each group. **F**. DCX-positive (immature), DCX-negative (mature), and average DCX-positive neuron soma volume data dimensionally reduced via a standard deviation scale to compare shifts across measures within and across individuals.

##### Mature (DCX-negative) neuron count estimates (Fig. 4B)

There was an average of 105,149 DCX-negative neurons present in infant monkeys (SD=37,948.33), ranging from 58,988 neurons - 147,633 neurons, and an average of 207,353 DCX-negative neurons (SD=36,672.27) present in adolescent monkeys, ranging from 168,454 neurons – 250,968 neurons, with a significant difference between groups (p=0.004). The number of putative mature neurons in the adolescent was approximately 197% of that in the 3 month-old infant PL, indicating an almost two-fold increase over the infant group.

We calculated the average sum of immature and mature neurons, and found that the infant PL average was 1,012,158 total neurons (SD=207,636.23) and the adolescent PL average was 748,518 total neurons (SD=68,848.82). There was a significant difference in total neuron numbers between groups (p=0.026), indicating decreasing total neuron numbers in the PL with age (Fig. 4C). Mean immature neuron numbers declined by 365,843, while mean mature neuron numbers increased by 102,204 between infancy and adolescence. Thus, declines in immature neurons were not completely offset by increases in mature neuron numbers in the PL by adolescence.

We then calculated the ratio of immature neurons per mature neuron in each group. Adolescent monkeys had a significantly smaller ratio of immature:mature neurons (on average 3 DCX-positive immature neurons for every 1 mature neuron) compared to infant monkeys (on average 9 DCX-positive immature neurons for every 1 mature neuron) (p = 0.003) (Fig. 4D). Both summed data (total DCX-positive plus DCX-negative neurons) and ratios of immature and mature neurons counts at infant and adolescent time points suggests several possibilities, including migration of neurons from the PL in the transition to adolescence (see Discussion).

##### Immature (DCX-positive) average neuron soma volume (Fig. 4E & Fig.5)

DCX-positive neuron soma volumes were measured as one feature of neuronal maturity, since increased DCX-positive soma volume is associated with a more mature neuronal state (Chareyron et al., 2012; Dalva, Ghosh, & Shatz, 1994; Kempermann, Jessberger, Steiner, & Kronenberg, 2004; Kohler, Williams, Stanton, Cameron, & Greenough, 2011). There was a significant difference in the average soma volume of DCX-positive cells in the PL of the infant monkey group (mean=190.75 μm^3^, SD=24.74, ranging from 164.48 μm^3^ – 220.45 μm^3^), and in the PL of the adolescent monkey group (mean=235.01 μm^3^, SD=30.87 ranging from 188.87 μm^3^ – 253.45 μm^3^) (Fig. 4E). This translates to an average DCX-positive neuron diameter of 7.14 μm in infants and 7.66 μm in adolescents. On average, DCX-positive neurons are significantly larger in adolescent monkeys compared to infant monkeys, (p=0.033), with an average DCX-positive soma volume increase of 7% compared to DCX-positive soma volume in infants.

**Figure 5:**
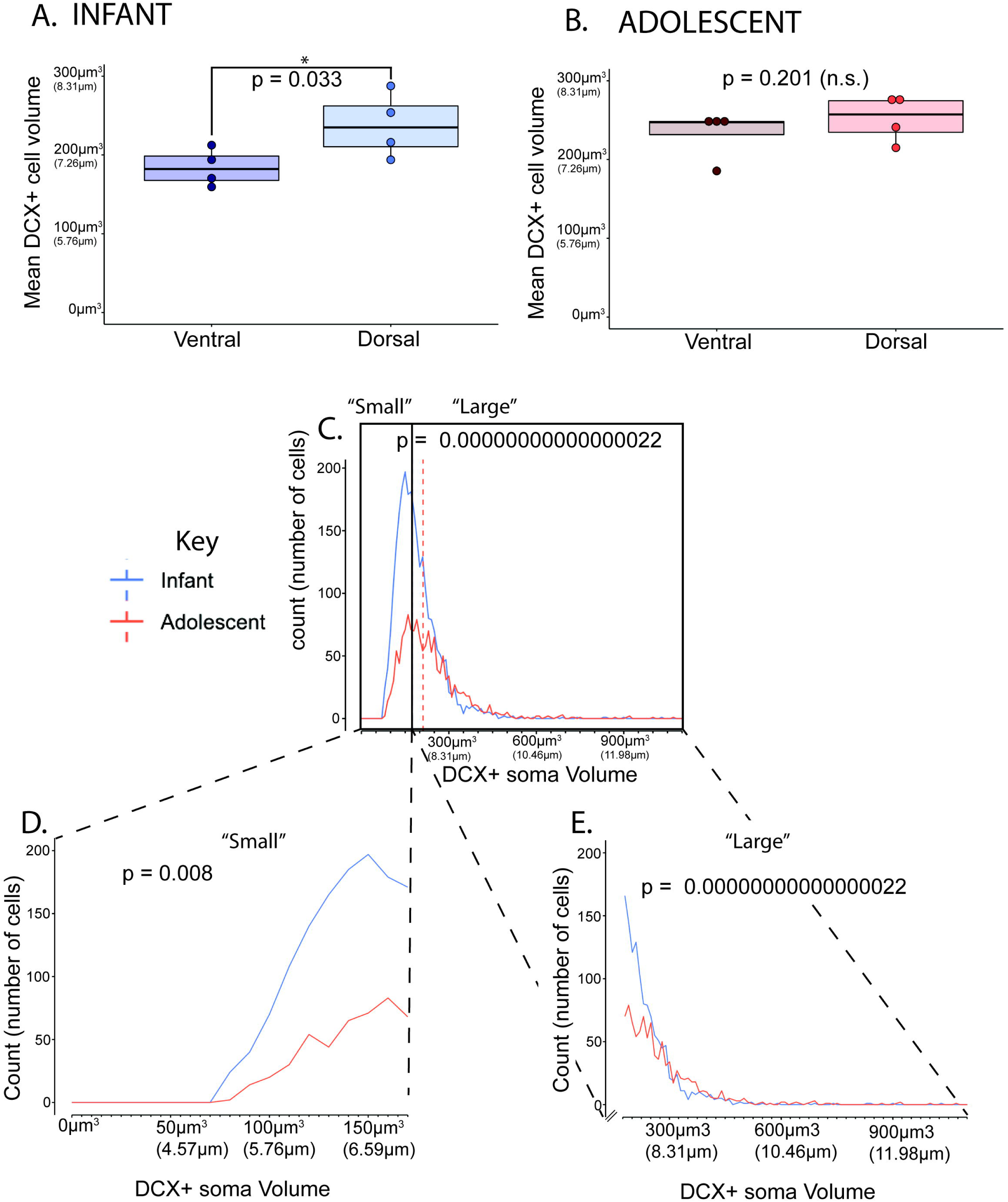
Normal Postnatal Development study: DCX-positive soma volumes. **A.** Boxplots showing infant control ventral and dorsal DCX-positive soma volumes (blue), and **B.** adolescent ventral and dorsal DCX-positive soma volumes. **C-E.** Frequency polygon plots showing (**C.)** all neurons with DCX-positive soma volumes measured ranging from 76.787 μm^3^ – 1067.568 μm^3^ (diameters of about 5.27 μm – 12.68 μm). Lines indicate the median DCX-positive soma volume of infant (blue) and adolescent (dotted orange) data (solid black line=median between “Small” and “Large” cells), (**D.)** “Small” DCX-positive soma volume neurons (174.38 μm^3^ and lower) with volumes ranging from 76.787 μm^3^ – 174.378 μm^3^ (diameters of about 5.27 μm – 6.93 μm), and (**E.)** “large” DCX-positive soma volume neurons (larger than 174.38 μm^3^), with volumes ranging from 174.396 μm^3^ – 1067.568 μm^3^ (diameters of about 6.93 μm – 12.68 μm).

Due to a qualitative difference between the dorsal-ventral gradient of DCX-positive soma volumes (Fig. 3), we examined for quantitative differences in dorsal-ventral DCX-positive soma volume for each animal. Here, we divided the PL into a dorsal 1/3 and a ventral 2/3. In infant control animals, DCX-positive soma volumes in the dorsal PL were significantly larger than DCX-positive soma volumes in the ventral PL, consistent with ventral to dorsal growth gradient in infants (dorsal soma volume average = 237.852 μm^3^ (7.687 μm), ventral soma volume average = 184.073 μm^3^ (7.058 μm), Shapiro-Wilk p = 0.600, Bartlett’s p = 0.385, one-tailed t-test p = 0.033, Cohen’s d = 1.592) (Fig. 5A). However, in adolescent animals, there was not a significant difference when comparing DCX-positive soma volumes between the dorsal PL and ventral PL (dorsal soma volume average = 252.021 μm^3^ (7.837 μm), ventral soma volume average = 232.503 μm^3^ (7.629 μm), Shapiro-Wilk p = 0.410, Bartlett’s p = 0.933, one-tailed t-test p = 0.201, Cohen’s d = 0.638) (Fig. 5B).

To examine DCX-positive soma volumes in the infant and adolescent PL samples as a whole, we first generated a frequency polygon plot containing the entire range of neuron DCX-positive soma volumes sampled (Fig. 5C, Table 2). A left-skewed distribution existed for DCX-positive soma volumes in both infant and adolescent PL, with small numbers of neurons with large DCX-positive soma volumes in each group, reflecting morphologic heterogeneity. Mean DCX-positive soma volumes between infant and adolescent control PL showed statistically significant differences such that mean DCX-positive soma volumes for adolescents were significantly larger than in infants (p = 0.00000000000000022)(Table 2).

**Table 2.**
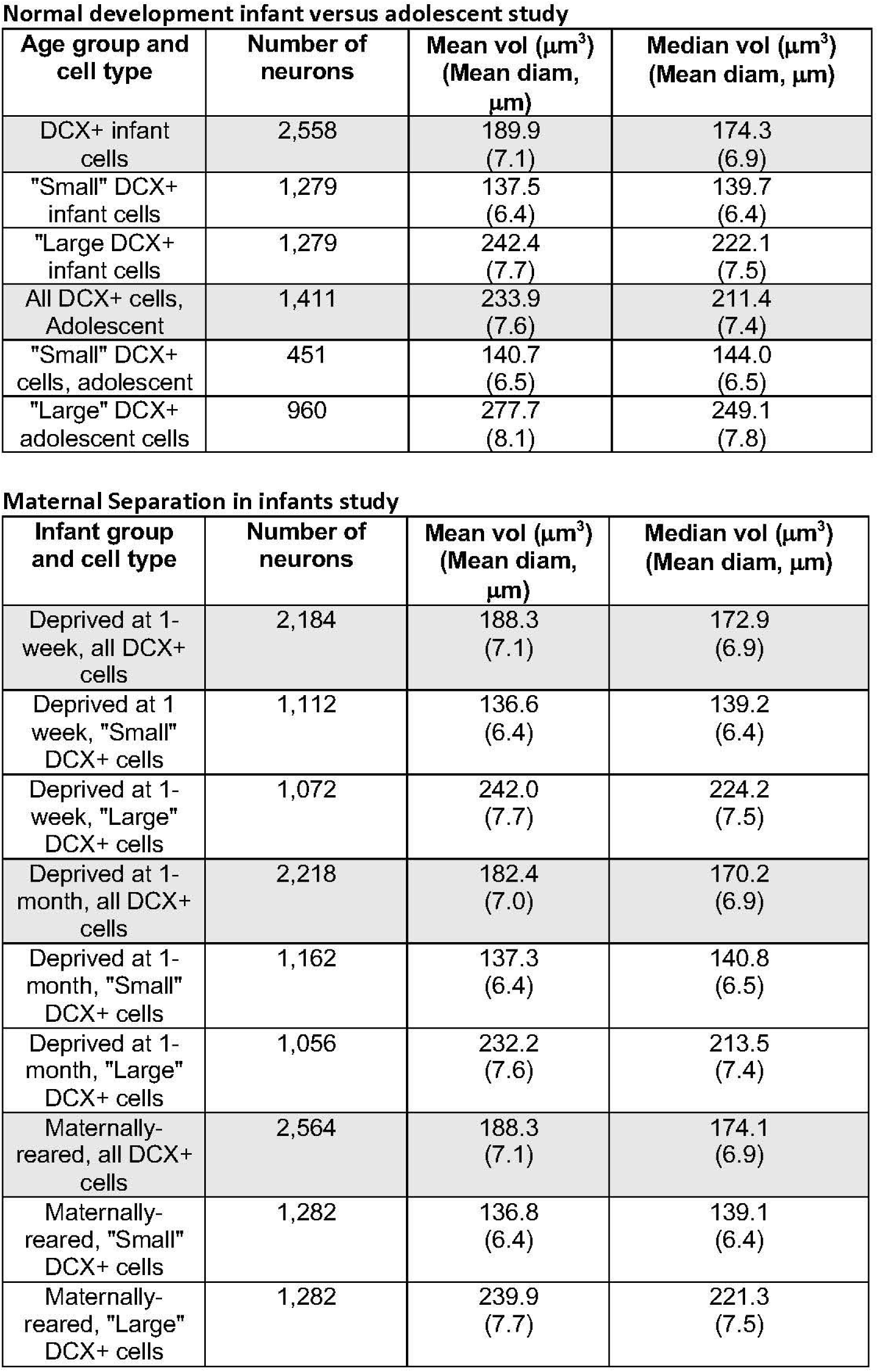
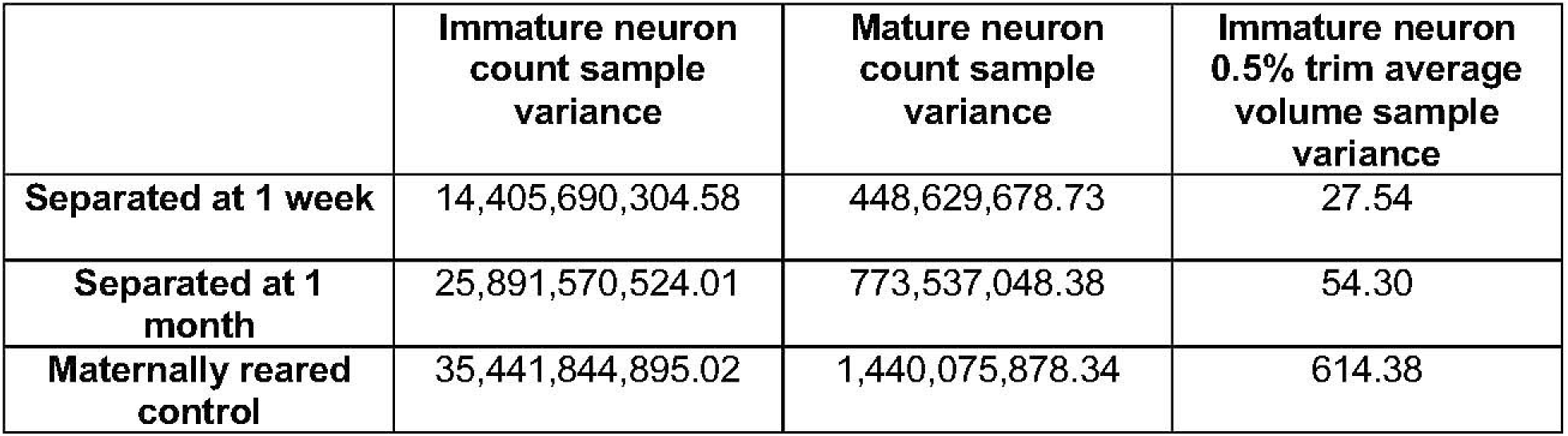
Soma volume means and medians.

**Normal development infant versus adolescent study**
| **Age group and cell type** | **Number of neurons** | **Mean vol ($\mu\text{m}^3$)**<br>**(Mean diam, $\mu\text{m}$)** | **Median vol ($\mu\text{m}^3$)**<br>**(Mean diam, $\mu\text{m}$)** |
| --- | --- | --- | --- |
| DCX+ infant cells | 2,558 | 189.9<br>(7.1) | 174.3<br>(6.9) |
| "Small" DCX+ infant cells | 1,279 | 137.5<br>(6.4) | 139.7<br>(6.4) |
| "Large" DCX+ infant cells | 1,279 | 242.4<br>(7.7) | 222.1<br>(7.5) |
| All DCX+ cells, Adolescent | 1,411 | 233.9<br>(7.6) | 211.4<br>(7.4) |
| "Small" DCX+ cells, adolescent | 451 | 140.7<br>(6.5) | 144.0<br>(6.5) |
| "Large" DCX+ adolescent cells | 960 | 277.7<br>(8.1) | 249.1<br>(7.8) |

**Maternal Separation in infants study**
| **Infant group and cell type** | **Number of neurons** | **Mean vol ($\mu\text{m}^3$)**<br>**(Mean diam, $\mu\text{m}$)** | **Median vol ($\mu\text{m}^3$)**<br>**(Mean diam, $\mu\text{m}$)** |
| --- | --- | --- | --- |
| Deprived at 1-week, all DCX+ cells | 2,184 | 188.3<br>(7.1) | 172.9<br>(6.9) |
| Deprived at 1 week, "Small" DCX+ cells | 1,112 | 136.6<br>(6.4) | 139.2<br>(6.4) |
| Deprived at 1-week, "Large" DCX+ cells | 1,072 | 242.0<br>(7.7) | 224.2<br>(7.5) |
| Deprived at 1-month, all DCX+ cells | 2,218 | 182.4<br>(7.0) | 170.2<br>(6.9) |
| Deprived at 1-month, "Small" DCX+ cells | 1,162 | 137.3<br>(6.4) | 140.8<br>(6.5) |
| Deprived at 1-month, "Large" DCX+ cells | 1,056 | 232.2<br>(7.6) | 213.5<br>(7.4) |
| Maternally-reared, all DCX+ cells | 2,564 | 188.3<br>(7.1) | 174.1<br>(6.9) |
| Maternally-reared, "Small" DCX+ cells | 1,282 | 136.8<br>(6.4) | 139.1<br>(6.4) |
| Maternally-reared, "Large" DCX+ cells | 1,282 | 239.9<br>(7.7) | 221.3<br>(7.5) |

|  | **Immature neuron**<br>**count sample**<br>**variance** | **Mature neuron**<br>**count sample**<br>**variance** | **Immature neuron**<br>**0.5% trim average**<br>**volume sample**<br>**variance** |
| --- | --- | --- | --- |
| **Separated at 1 week** | 14,405,690,304.58 | 448,629,678.73 | 27.54 |
| **Separated at 1**<br>**month** | 25,891,570,524.01 | 773,537,048.38 | 54.30 |
| **Maternally reared**<br>**control** | 35,441,844,895.02 | 1,440,075,878.34 | 614.38 |

We calculated the median DCX-positive soma volume of the maternally-reared infants (DCX-positive soma volume = 174.38 μm^3^, equivalent to a diameter of 6.93 μm) as the boundary that distinguished between “small” and “large” neuron DCX-positive soma volumes for both infant and adolescent controls. Here we found that adolescent ‘small’ volume somas are significantly larger than infant ‘small’ volume somas (p = 0.008) (Fig. 5D), and also that adolescent ‘large’ volume somas are significantly larger than infant ‘large’ volume somas (p = 0.00000000000000022) (Fig. 5E, Table 2). Thus, despite significant heterogeneity in cell body sizes in each group, DCX-positive soma volumes are overall significantly larger in adolescent PL compared to infant P, suggesting overall DCX-positive soma size with age.

##### Between-subjects comparisons for normal development data (Fig. 4F)

In order to more clearly track between-subjects variation, we dimensionally reduced DCX-positive (immature) cell counts, DCX-negative (mature) cell counts, and average DCX-positive soma volume into standard deviation scales <u>to track relationships among “high” value of mature count, “low” value of immature count, and DCX-positive soma size for each individual in each group</u> (Fig. 4F).

Overall, there were decreased numbers of immature neurons, increased numbers of mature neurons, and increased DCX-positive immature neuron soma volumes between infant and adolescent cohorts, in the face of a generally decreasing total neuron population in the PL.

### Maternal separation study in infant macaques (STUDY 2)

#### Cellular measures in maternally-separated and maternally-reared infants

Because there were pronounced changes in neuron numbers and DCX-positive immature neuron soma volume between infant and adolescent control animals, we examined if early-life environmental perturbations in the form of maternal separation led to effects on these same variables in infants that were maternally-reared or maternally-separated. The same maternally-reared (control) infant group data used in the normal development study (above) were compared to the same measures collected in 2 maternally-separated infant groups (infants separated from their mother at 1-week of age and infants separated from their mother at 1-month of age).

##### Immature (DCX-positive) neuron count estimates (Fig. 6A)

There were no significant differences in the average number of DCX-positive neurons between separated at 1-week, separated at 1-month, and maternally-reared control groups (F_(2,9)_ = 0.728, p = 0.509. Separated at 1-week data: average = 917,419 neurons, range = 802,352 – 1,025,374 neurons, SD 120,023.71; Separated at 1-month data: average = 795,138 neurons, range = 624,574 – 996,134 neurons; SD = 160,908.58; Control data: average = 907,008 neurons, range = 678,985 – 1,129,609 neurons, SD = 188,260.05).

**Figure 6:**
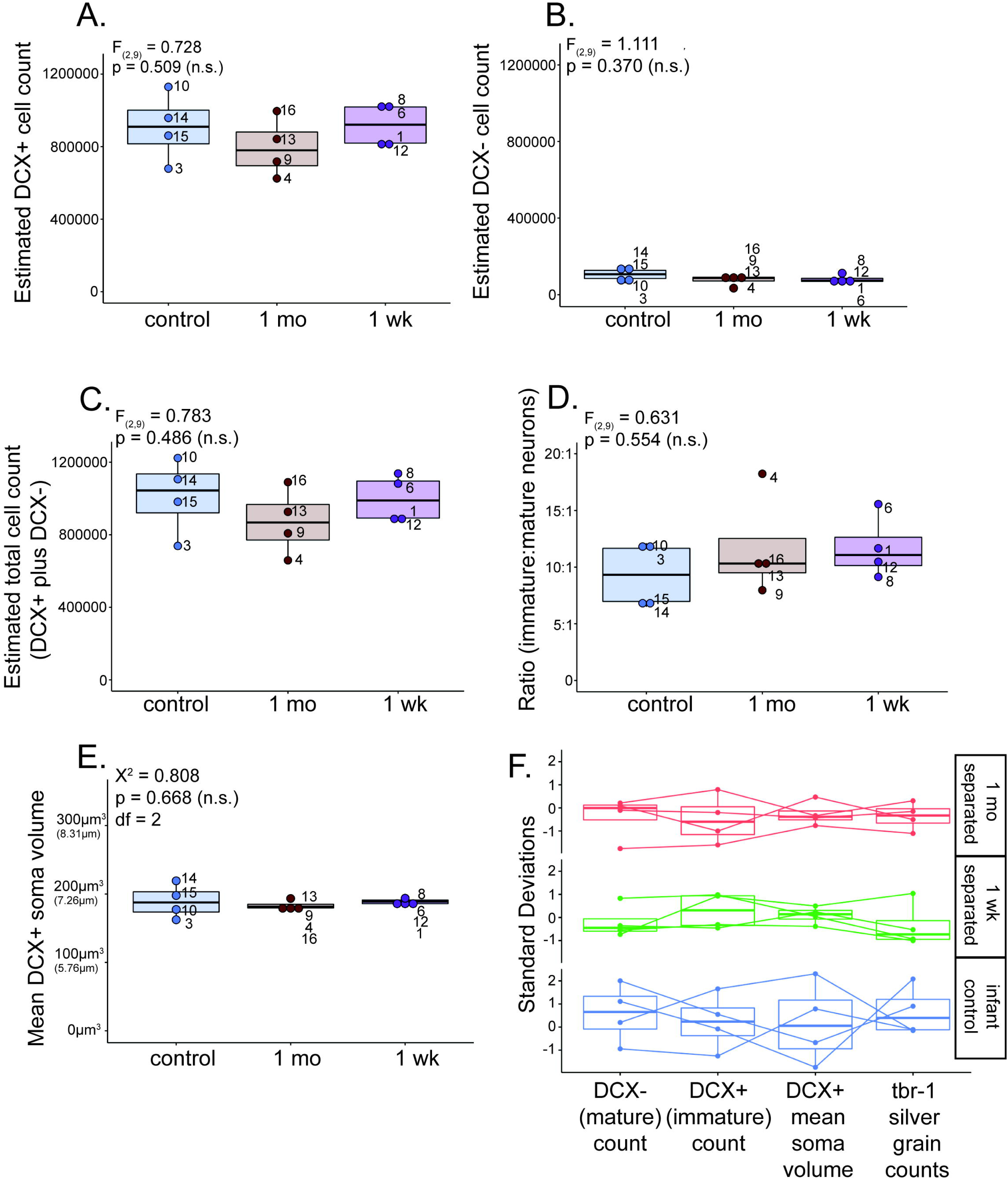
Maternal separation study: Cellular measurements. **A-E.** Boxplots showing maternally-reared infant control (blue), separated at 1-month (brown), and separated at 1-week (magenta), **A.** DCX-positive (immature) neuron count estimate data, **B.** DCX-negative (mature) neuron count estimate data, **C.** Summed DCX-positive (immature) and DCX-negative (mature) neuron counts, **D.** Ratio of DCX-positive (immature) to DCX-negative (mature) neuron counts, and **E.** average DCX-positive neuron soma volume data. Small numbers listed next to data points represent identification numbers for individual animals in each group. **F**. DCX-positive (immature), DCX-negative (mature), average DCX-positive neuron soma volume, and *tbr-1* mRNA levels data <u>dimensionally reduced via a standard deviation scale to directly compare how each of these variables shift within and across individuals.</u>

##### Mature (DCX-negative) neuron count estimates (Fig. 6B)

No significant differences were found in the average number of DCX-negative neurons across separated at 1-week, separated at 1-month, and maternally-reared control groups (F_(2,9)_ = 1.111, p = 0.370. Separated at 1-week data: average = 81,261 neurons, range = 65,342 – 112,270 neurons, SD = 21,180.88; Separated at 1-month data: average = 75,550 neurons, range = 34,252 – 93,692 neurons, SD=27,812.53; control data: average = 105,149 neurons, range = 58,988 neurons – 147,633 neurons, SD = 37,948.33).

We calculated the average sum of immature and mature neurons for maternally-separated and infant control groups (Fig. 6C). The average sum of immature and mature neurons in separated at 1-week animals was 998,680 neurons (SD = 130500.40), in separated at 1-month animals was 870,687 neurons (SD = 182508.93), and in infant control animals was 1,012,158 neurons (SD = 207636.23). There was no significant difference in total neuron numbers between groups (F_(2,9)_ = 0.783, p = 0.486), indicating that total neuron numbers in the infant PL do not change in response to early-life maternal separation by 3 months of age. Furthermore, we calculated the ratio of immature:mature neurons, and found that there were no significant differences in immature:mature neuron ratios across maternally deprived and control infant monkey groups (F_(2,9)_ = 0.631; p = 0.554) (Fig. 6D).

##### Immature (DCX-positive) soma volume (Fig. 6E, & Fig. 7)

No significant differences were found for the average DCX-positive soma volume in the whole PL across separated at 1-week, separated at 1-month, and maternally-reared control groups (χ^2^ = 0.808, df = 2, p = 0.668. Separated at 1-week data: average = 188.24 μm^3^, range = 181.43 – 193.84 μm^3^, SD = 5.25; Separated at 1-month data: average = 183.10 μm^3^, range = 176.10 – 193.46 μm^3^, SD = 7.37; control data: average = 189.26 μm^3^, range = 162.37 – 219.93 μm^3^, SD = 24.79) (Fig. 6E). These average DCX-positive soma volumes are equivalent to diameters of 7.11 μm (separated at 1-week), 7.05 μm (separated at 1-month), and 7.12 μm (maternally-reared control).

**Figure 7:**
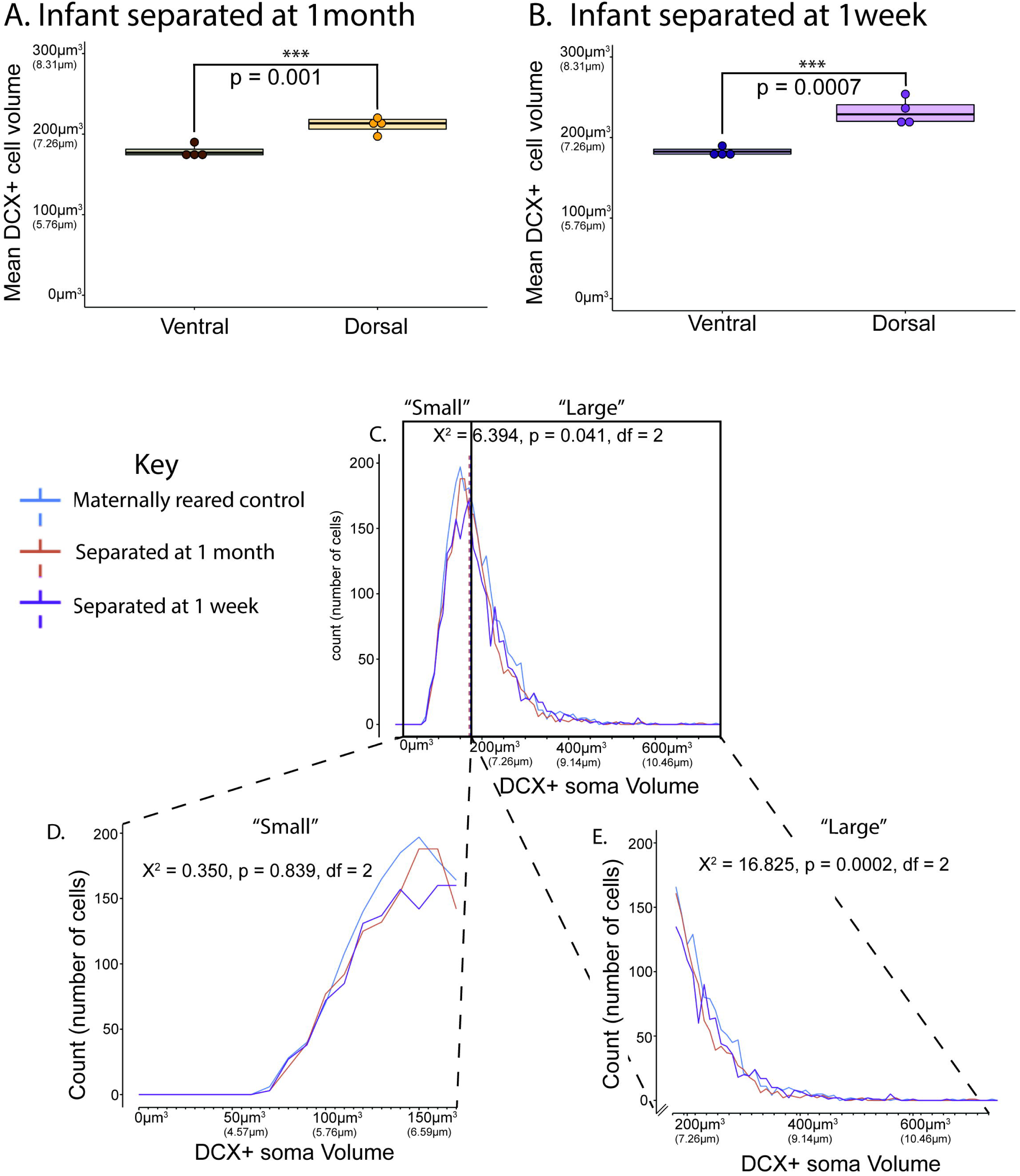
Maternal separation study: DCX-positive soma volume. **A-B.** Boxplots showing dorsal versus ventral mean soma volumes for **A.)** separated at 1-month (brown) and **B.)** separated at 1-week (purple) groups. Infant maternally-reared control shown in Fig. 5A. **C-E.** Frequency polygon plots showing **C.** all neurons with DCX-positive soma volumes measured ranging from 72.654 μm^3^ – 742.323 μm^3^ (a diameter of about 5.18 μm – 11.23 μm). Line indicates the median DCX-positive soma volume of separated at 1-month (dotted yellow), separated at 1-week (dotted magenta) and maternally-reared infant (blue). **D.** “Small” DCX-positive soma volume neurons (174.09 μm^3^ and lower) with volumes ranging from 72.654 μm^3^ – 174.081 μm^3^ (diameters of about 5.18 μm – 6.93 μm), and **E.** “large” DCX-positive soma volume neurons (larger than 174.09 μm^3^) ranging from 174.092 μm^3^ – 742.323 μm^3^ (diameters of about 6.93 μm – 11.23 μm).

DCX-positive soma volumes were significantly larger in dorsal PL compared to ventral PL for both groups of separated animals ((Fig. 7A-B). Infants separated beginning at 1 month (dorsal soma volume average = 211.034 μm^3^ (7.387 μm), ventral soma volume average = 178.618 μm^3^ (6.987 μm), Shapiro-Wilk p = 0.413, Bartlett’s p = 0.714, one-tailed t-test p = 0.001, Cohen’s d = 3.464) (Fig. 7A) and infants separated beginning at 1 week (dorsal soma volume average = 232.219 μm^3^ (7.626 μm), ventral soma volume average = 182.380 μm^3^ (7.036 μm), Shapiro-Wilk p = 0.3747, Bartlett’s p = 0.1424, one-tailed t-test p = 0.0007, Cohen’s d = 3.968) showed significant differences in between these regions (Fig. 7B). Overall, for all monkey groups examined (including normal control infants, Fig. 5A), dorsal soma volume is generally larger than “whole” soma volume average, with infant groups having slightly smaller ventral soma volume average compared to “whole” soma volume average.

Frequency polygon plots of the distribution of DCX-positive soma volumes in the PL between separated at 1-week, separated at 1-month, and maternally-reared control animals (Fig. 7C-E, Table 2) showed significant differences among groups (χ^2^ = 6.394, df = 2, p = 0.041), with separated at 1-month animals vs maternally-reared animals driving this difference (Dunn’s test with BH correction: separated at 1-month vs. separated at 1-week p= 0.083; separated at 1-month vs. maternally-reared p= 0.049, separated at 1-week vs. maternally-reared p= 0.683) (Fig. 7C).

After sorting DCX-positive neurons into “small” and “large” subpopulations across groups, using the median DCX-positive soma volume from the maternally-reared infant group as a break point (median DCX-positive soma volume=174.09 μm^3^, equivalent to a diameter of 6.93 μm) (Fig. 7D-E, Table 2), we found no significant differences between groups when comparing small neurons (χ^2^ = 0.350, df = 2, p = 0.839) (Fig. 7D). However, large neurons in the separated at 1-month group had significantly smaller volumes compared to both maternally-reared and separated at 1-week groups (χ^2^ = 16.825, df = 2, p = 0.0002; Dunn’s test with BH corrections: separated at 1-month vs. separated at 1-week p = 0.0005; separated at 1-month vs. maternally-reared p = 0.001, separated at 1-week vs. maternally-reared p = 0.567) (Fig. 7E).

##### Between-subjects comparisons for maternal separation data (Fig. 6F)

To examine between-subjects variation, we dimensionally reduced DCX-positive (immature) cell counts, DCX-negative (mature) cell counts, average DCX-positive soma volume, and *tbr-1* mRNA levels to examine standard deviation in each measure for each animal across group means (Fig. 6F).

Testing for equality of variance revealed the variability of average immature neuron soma volume across infant groups, which was significantly larger for maternally-reared infants compared to both separation groups (χ^2^ = 6.821, p = 0.033). This difference was not significant for immature (χ^2^ = 0.514, p = 0.774) and mature (χ^2^ = 0.876, p = 0.645) neuron counts. Reduced variability in soma volume in separated infants, or greater clustering around the mean, may suggest that separation had a general ‘braking’ effect on individual immature neuron growth. This would be consistent with our previous finding that maternal separation-induces reductions in PL tbr1 mRNA—a neuronal growth promotor--in separated infants (D. M. de Campo et al., 2017) (see below).

#### Correlations among immature and mature neuron number, cell soma size, and tbr-1 levels in infant groups

Correlation analyses were conducted in the infant maternal separation study because correlations across groups can capture biological processes that change together when an environmental stressor is introduced. Moreover, individual differences in stress sensitivity of biologic systems can be visualized (Flagel, Akil, & Robinson, 2009). In this section, we examined correlations across cellular measurements we obtained (i.e. correlating DCX-positive immature soma volume with immature neuron counts and mature neuron counts), and additionally examined correlations between *tbr-1* levels and cellular measurements obtained (i.e. estimated number of immature neurons, estimated number of mature neurons, and average DCX-positive immature neuron soma volume) from the same cohort.

##### Correlations among PL cellular measurements (Fig. 8A)

To determine if DCX-positive immature neuron soma volume predicted numbers for either immature neurons or mature neurons, we first conducted one-tailed Pearson’s correlations between average DCX-positive immature soma volume vs. immature neuron counts, and average DCX-positive immature neuron soma volume vs. mature neuron counts. There was a significant correlation between average DCX-positive immature soma volume vs. mature neuron counts (r=0.724, p=0.004), but no significant correlation between average DCX-positive immature soma volume vs. immature neuron counts (r=0.312, p=0.162). These findings indicate that increased DCX-positive soma volume tracks mature neuron numbers across all infants.

**Figure 8:**
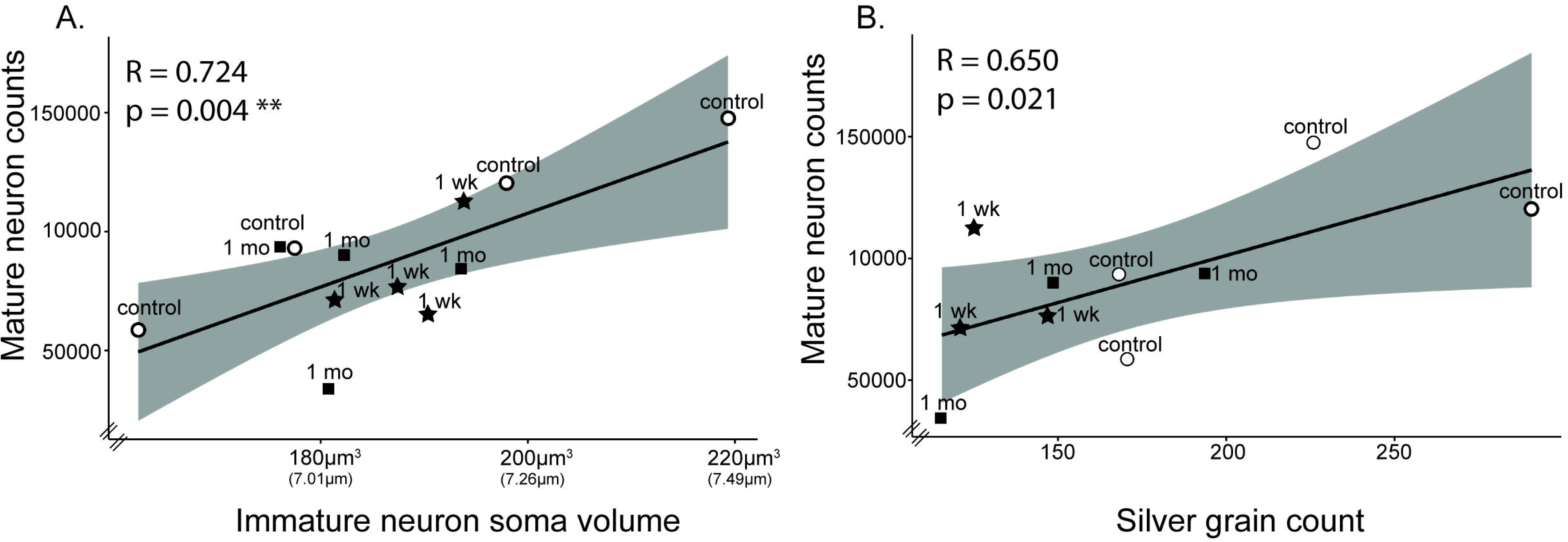
Maternal separation study: Correlations between DCX-positive immature neuron soma volume, mature neuron count, and tbr-1 levels. **A.** DCX-positive immature neuron soma volume correlates vs. mature neuron counts. **B.** *tbr-1* mRNA levels vs. DCX-negative (mature) neuron count estimates. Maternally-reared (control) datapoints are indicated with white circles and abbreviated “control”, separated beginning at 1-month data points are indicated with black squares and abbreviated “1 mo”, and separated beginning at 1 week data points are indicated with black stars and abbreviated “1 wk”. Gray shading indicates 95% confidence intervals.

##### Correlations between cellular measurements and tbr-1 levels (Fig. 8B)

*tbr-1* mRNA is an activity dependent molecule that is required for neural growth (S. F. Darbandi et al., 2020; Mihalas & Hevner, 2017). We previously found that it is significantly decreased in the PL after 1-week and 1-month maternal separation in this cohort (D. M. de Campo et al., 2017). One goal in quantifying cellular measures was to determine whether *tbr-1* mRNA levels were associated with indices of cellular maturation. We correlated *tbr-1* mRNA levels in each animal with immature or mature neuron counts, and immature neuron soma size. There was a significant positive correlation between *tbr-1* mRNA levels vs. mature neuron counts (r=0.650, p=0.021), but no significant correlations between *tbr-1* mRNA levels vs. immature neuron counts (r=0.245, p=0.247), or between *tbr-1* mRNA levels vs. average DCX-positive immature soma volume (r=0.443, p=0.086). Thus, *tbr-1* mRNA levels specifically predict increases in mature neuron numbers, defined by lack of DCX-protein. This is consistent with *tbr-1’s* role in the late stages of immature glutamatergic neuron differentiation and connectivity (Siavash Fazel Darbandi et al., 2018).

## Discussion

### Summary of findings

The PL is a little understood region of the amygdala, and even less is known about how it changes in response to normal postnatal development or early life stress. In the context of normal postnatal development, we found that immature neuron numbers decrease, and mature neuron numbers increase, over a backdrop of decreasing total neuron numbers between female infant and adolescent time points. DCX-positive immature neuron soma volume is consistently larger in adolescent compared to infant monkeys, even when sub-classifying neurons into “small” and “large” volume categories. These findings replicate and extend prior studies (Chareyron et al., 2012) and provide strong evidence that in normal young female macaques, PL neurons mature between infancy and adolescence. Our finding that total DCX-positive and DCX-negative neurons decrease in PL neuron number between infancy and adolescence is consistent with recent findings (Chareyron, Banta Lavenex, Amaral, & Lavenex, 2021), and implies neuron migration or removal from the PL over early development.

In the infant female maternal separation study, there were no significant differences between separated and maternally-reared animals in immature or mature cell numbers by 3 months of age. Yet, we previously found that maternal separation is associated with significant decreases in several transcripts associated with neural differentiation (D. M. de Campo et al., 2017). The fact that molecular changes did not translate to group differences in neuronal numbers by 3 months of age is likely due to the short interval between the maternal separation and sacrifice (1-2 months). Cellular maturation occurs over a protracted, months-long period in nonhuman primates (Kohler et al., 2011). Examination of the effects of maternal separation on immature and mature cell numbers in somewhat older infant monkeys may be more informative.

Despite the lack of overall differences in immature and mature cellular numbers, some shifts in soma volume variability were detected in separated infants. The variance in all soma volume was much larger for control infant PL neurons compared to that in separated groups, suggesting that normal variation in cellular growth is damped by maternal separation, leading to more “homogeneous” cellular fates. In addition, infants that were maternally-separated at 1-month had significantly smaller DCX-positive “large” neurons, suggesting a possible “stunting” of soma size enlargement in this group.

Investigating correlations at the individual level across the infant groups, we found that DCX-positive soma volume and mature neuron counts were positively correlated with one another, while DCX-positive soma volume and immature neuron counts were not, in-line with our predictions. *tbr-1* mRNA levels also positively correlated with increased neural maturation, suggesting that higher *tbr-1* mRNA levels are associated with overall greater neuronal maturity. Despite individual variability, maternally separated animals tended to have lower numbers of mature neurons and lower tbr-1 mRNA levels in the PL.

### Evidence for neurons maturing and migrating with age in the PL

Previous studies indicate that between infancy and adulthood, mature neuron numbers and neuron size increase in the PL in nonhuman primate (Chareyron et al., 2012), and in humans (Avino et al., 2018). Post-mortem work in human shows that immature PL neurons in young brains gradually transition into mature, glutamatergic neurons (staining positively for *tbr-1* and VGLUT2) (Sorrells et al., 2019). Consistent with these data, we found that the ratio of immature to mature neuron numbers decreases between infancy and adolescence but not when comparing maternally separated infant groups to a control infant group. We additionally found that total DCX-positive (immature) and DCX-negative (mature) neurons decrease in the PL between infant and adolescent cohorts. This results conflicts with previous results using Nissl-staining identification of neuron types, and different age-groups, which found no change (Chareyron et al., 2012). Use of DCX-immunostaining for PL definition and counting, as well as less age homogeneity in the adolescent group (4 years here versus 5-9 years in Chareyron et al.) may explain some of these differences. In more recent work, Chareyron et al. also did not find changes in total PL neuron counts when comparing infant monkey groups aged 0-3 months, 6-9 months, and 1 year old, although this time period is much shorter than that in the present study (Chareyron et al., 2021).

Reasons for the decline in total neuron numbers cannot be answered from our study. While cell loss between infancy and adolescence may be due to apoptosis, we found few apoptotic profiles (pyknotic cells) in either sample. However, it is still possible that the absence of pyknotic nuclei could be explained by rapid clearance of cells by microglia. However, it is also likely that immature neurons are maturing and migrating out of the PL, with the basal nucleus acting as a putative target region for newly mature neurons exiting the PL. This would be consistent with the ‘gradient of soma’ size found in our study, and the presence of scattered large DCX-positive neurons in the Bpc. Recent human studies also support the idea of a ‘dorsal-ward’ migration, since mature neurons increase in the overlying basal nucleus by 30% between infancy and early life, while immature PL neurons decline (Avino et al., 2018). Together, these findings lend further evidence to the idea that the PL houses a pool of smaller, newly mature neurons that undergo protracted maturation, possibly migrating into surrounding amygdala, or other brain regions, over development. We plan to assess basal nucleus neuron counts in future work.

### Molecular cascades in maternally-separated animals

Glutamatergic projection neuron development results from a cascading molecular process that unfolds over time (Hodge, Kahoud, & Hevner, 2012). *tbr-1* is expressed in post-mitotic neurons, overlapping and extending past the period of DCX-expression (Hevner, Hodge, Daza, & Englund, 2006). In development, *tbr-1* protein is induced by excitatory transmission, and plays roles in synaptogenesis (Siavash Fazel Darbandi et al., 2018; Huang et al., 2014), dendritic patterning, and laminar specification (Bedogni et al., 2010; Siavash Fazel Darbandi et al., 2018). These activities herald the transition of “immature” cells into functioning neurons that participate in circuits. In our previous work, we found that maternal separation at 1-week and 1-month results in reduction of PL *tbr-1* expression, and other molecules involved in post-mitotic neural growth: e.g. anaphase-promoting complex (*apc,* regulation of axonal elongation, synapse size and number), neural cell adhesion molecule 1 (*ncam1*, which regulates neurite outgrowth, cell migration, and neurogenesis), neuritin 1 (*nrn1*, which is associated with adult neuroplasticity and is found in differentiating, post-mitotic neurons) and a RAS-related protein (*rap1a*, a small GTPase associated with cell adhesion and proliferation) (D. M. de Campo et al., 2017).

In the present study, we found that that *tbr1* mRNA levels (measured using *in situ* hybridization) predict increased immature neuron soma size (associated with more mature phenotypes), and mature neuron numbers across individuals, consistent with its known functions in the developing brain. This unsurprising since *tbr1* expression drives a number of molecular cascades involved in neuronal maturation (Huang et al., 2014)(S. F. Darbandi et al., 2020), and soma volume is associated with a more mature neuron state(Chareyron et al., 2012; Kempermann et al., 2004; Kohler et al., 2011) (Purves, Snider, & Voyvodic, 1988). Having soma volumes increase in an age-appropriate manner may allow for more numerous synaptic connections with both local and global pathways, which can therefore integrate these larger neurons into a wider range of circuits important for proper amygdala functioning. Although speculative, since separated at 1-month animals had fewer “large” immature neurons, it may be that undergoing maternal deprivation at this developmental time point may shift the standard timeline for neuron soma volume enlargement, therefore processing speed and synaptic connections may not form optimally due to soma volumes (and accompanying dendrites and axons) being smaller than they should be at that given timepoint. Although not examined in the current study, maternal separation is associated with decreased pro-social behavior compared to maternally-reared control animals (Sabatini et al., 2007)(DeCampo et al, 2017), and misaligned timing of soma volume development in the PL that is associated with maternal separation may partially influence these social behavior deficits.

### Limitations and future directions

Although there was clear evidence of neuronal growth occurring in the PL when comparing infant and adolescent monkeys, evidence for neuronal changes in the PL across maternally-separated infants was more limited, even though the stress of early-life maternal separation has well-known adverse impacts on the amygdala and its circuitry (Gee et al., 2013; Sabatini et al., 2007; Tottenham et al., 2011). This lack of evidence may be due to the small sample sizes in the current study. Another possibility is that the presence of other females in our separation model, may buffer more severe brain effects. Finally, we speculate that scale changes in neuronal numbers may not be apparent in the two-month timeframe between the manipulation and sacrifice, given the length of neural development in primates (Kohler et al., 2011). On the other hand, early neuronal change is detected in molecular shifts in the PL, and also in the reduced variability among separated animals in soma size compared to maternally reared infants. One interpretation is that this represents a harbinger of decreased neural maturation that might be evident at later time-points. Future studies will examine larger samples, and older animals.

Another limitation is the lack of cortisol levels for our animal cohorts. Therefore, we are unable to directly link maternal separation brain changes to ‘stress’ effects. Indeed, brain changes may be due to decreased sensory/emotional inputs, immune factors, or other mechanisms.

## Conclusion

The PL is a unique brain region of immature neurons that changes over the course of postnatal development, and may be altered in response to environmental influences. The present study shows that there are significant cellular changes in female infant and adolescent monkeys; there are fewer immature neurons, larger immature neurons, and more mature neurons with age. Findings from our previous work showed that maternal separation significantly reduced *tbr-1* mRNA levels (DeCampo et al, 2017). In our current study, we examined cellular correlates, and found that both *tbr-1* mRNA levels and DCX-positive soma volume size correlated with mature neuron counts across all infant animals even though maternal separation did not result in clear differences in neuron counts across infant groups. Reduced variance in all cellular measures in separated infants, combined with reductions in tbr1 gene expression, may suggest early effects of maternal separation on cellular growth in these groups.

## Acknowledgements

This work was funded through the support of the National Institute of Mental Health R21MH127486 (J.L.F., J.C.), and the Schmitt Program on Integrative Neuroscience (SPIN) (J.L.F.).

## References

Amaral, D. G., & Bassett, J. L. (1989). Cholinergic innervation of the monkey amygdala: An immunohistochemical analysis with antisera to choline acetyltransferase. J. Comp. Neurol., 281, 337–361.

Avino, T. A., Barger, N., Vargas, M. V., Carlson, E. L., Amaral, D. G., Bauman, M. D., & Schumann, C. M. (2018). Neuron numbers increase in the human amygdala from birth to adulthood, but not in autism. Proc. Natl. Acad. Sci. U. S. A. doi:10.1073/pnas.1801912115

Bedogni, F., Hodge, R. D., Elsen, G. E., Nelson, B. R., Daza, R. A., Beyer, R. P., … Hevner, R. F. (2010). Tbr1 regulates regional and laminar identity of postmitotic neurons in developing neocortex. Proc Natl Acad Sci U S A, 107(29), 13129–13134. doi:10.1073/pnas.1002285107

Braak, H., & Braak, E. (1983). Neuronal types in the basolateral amygdaloid nuclei of man. Brain. Res. Bull., 11(3), 349–365.

Bulfone, A., Smiga, S. M., Shimamura, K., Peterson, A., Puelles, L., & Rubenstein, J. L. (1995). T-brain-1: a homolog of Brachyury whose expression defines molecularly distinct domains within the cerebral cortex. Neuron, 15(1), 63–78.

Cameron, J. L. (2001). Critical periods for social attachment: deprivation and neural systems in rhesus monkeys. Paper presented at the Soc Res Child Development.

Cameron, J. L., Eagleson, K. L., Fox, N. A., Hensch, T. K., & Levitt, P. (2017). Social Origins of Developmental Risk for Mental and Physical Illness. J. Neurosci., 37(45), 10783–10791. doi:10.1523/JNEUROSCI.1822-17.2017

Chareyron, L. J., Amaral, D. G., & Lavenex, P. (2016). Selective lesion of the hippocampus increases the differentiation of immature neurons in the monkey amygdala. Proc. Natl. Acad. Sci. U. S. A., 113(50), 14420–14425. doi:10.1073/pnas.1604288113

Chareyron, L. J., Banta Lavenex, P., Amaral, D. G., & Lavenex, P. (2021). Life and Death of Immature Neurons in the Juvenile and Adult Primate Amygdala. Int J Mol Sci, 22(13). doi:10.3390/ijms22136691

Chareyron, L. J., Lavenex, P. B., Amaral, D. G., & Lavenex, P. (2012). Postnatal development of the amygdala: A stereological study in macaque monkeys. J. Comp. Neurol., 520(9), 1965–1984. doi:10.1002/cne.23023

Dalva, M. B., Ghosh, A., & Shatz, C. J. (1994). Independent control of dendritic and axonal form in the developing lateral geniculate nucleus. J Neurosci, 14(6), 3588–3602.

Darbandi, S. F., Schwartz, S. E. R., Pai, E. L. L., Everitt, A., Turner, M. L., Cheyette, B. N. R., … Rubenstein, J. L. R. (2020). Enhancing WNT Signaling Restores Cortical Neuronal Spine Maturation and Synaptogenesis in Tbr1 Mutants. Cell. Rep., 31(2), 831–845. doi:ARTN 107495 10.1016/j.celrep.2020.03.059

Darbandi, S. F., Schwartz, S. E. R., Qi, Q., State, M. W., Sohal, V. S., & Rubenstein, J. L. R. (2018). Neonatal Tbr1 Dosage Controls Cortical Layer 6 Connectivity. Neuron, 100(November 21, 2018), 831–845. doi:https://doi.org/10.1016/j.neuron.2018.09.027

de Campo, D., & Fudge, J. L. (2011). Where and what is the paralaminar nucleus? A review on a unique and frequently overlooked area of the primate amygdala. Neurosci Biobehav Rev, in revision.

de Campo, D. M., Cameron, J. L., Miano, J. M., Lewis, D. A., Mirnics, K., & Fudge, J. L. (2017). Maternal deprivation alters expression of neural maturation gene tbr1 in the amygdala paralaminar nucleus in infant female macaques. Dev. Psychobiol., 59(2), 235–249. doi:10.1002/dev.21493

Flagel, S. B., Akil, H., & Robinson, T. E. (2009). Individual differences in the attribution of incentive salience to reward-related cues: Implications for addiction. Neuropharmacology, 56 Suppl 1, 139–148. doi:10.1016/j.neuropharm.2008.06.027

Fudge, J. L. (2004). Bcl-2 immunoreactive neurons are differentially distributed in subregions of the amygdala and hippocampus of the adult macaque. Neuroscience, 127, 539–556.

Fudge, J. L., deCampo, D. M., & Becoats, K. T. (2012). Revisiting the hippocampal-amygdala pathway in primates: association with immature-appearing neurons. Neuroscience, 212, 104–119. doi:10.1016/j.neuroscience.2012.03.040

Garcia-Cabezas, M. A., John, Y. J., Barbas, H., & Zikopoulos, B. (2016). Distinction of Neurons, Glia and Endothelial Cells in the Cerebral Cortex: An Algorithm Based on Cytological Features. Front Neuroanat, 10, 107. doi:10.3389/fnana.2016.00107

Gee, D. G., Gabard-Durnam, L. J., Flannery, J., Goff, B., Humphreys, K. L., Telzer, E. H., … Tottenham, N. (2013). Early developmental emergence of human amygdala-prefrontal connectivity after maternal deprivation. Proc. Natl. Acad. Sci. U. S. A. doi:10.1073/pnas.1307893110

Geneser-Jensen, F. A., & Blackstad, T. W. (1971). Distribution of acetyl cholinesterase in the hippocampal region of the guinea pig. Z. Zellforsch, 114, 460–481.

Gundersen, H. J. (1988). The nucleator. J Microsc, 151(Pt 1), 3–21. doi:10.1111/j.1365-2818.1988.tb04609.x

Gundersen, H. J. G., & Jensen, E. B. (1987). The Efficiency of Systematic-Sampling in Stereology and Its Prediction. Journal of Microscopy-Oxford, 147, 229–263. doi:Doi 10.1111/J.1365-2818.1987.Tb02837.X

Harlow, H. F., Dodsworth, R. O., & Harlow, M. K. (1965). Total social isolation in monkeys. Proc. Natl. Acad. Sci. U. S. A., 54(1), 90–97.

Hevner, R. F., Hodge, R. D., Daza, R. A., & Englund, C. (2006). Transcription factors in glutamatergic neurogenesis: conserved programs in neocortex, cerebellum, and adult hippocampus. Neurosci. Res., 55(3), 223–233. doi:10.1016/j.neures.2006.03.004

Hevner, R. F., Shi, L., Justice, N., Hsueh, Y., Sheng, M., Smiga, S., … Rubenstein, J. L. (2001). Tbr1 regulates differentiation of the preplate and layer 6. Neuron, 29(2), 353–366.

Hinde, R. A., Spencer-Booth, Y., & Bruce, M. (1966). Effects of 6-day maternal deprivation on rhesus monkey infants. Nature, 210(5040), 1021–1023.

Hodge, R. D., Kahoud, R. J., & Hevner, R. F. (2012). Transcriptional control of glutamatergic differentiation during adult neurogenesis. Cellular and molecular life sciences : CMLS, 69(13), 2125–2134. doi:10.1007/s00018-011-0916-y

Howell, B. R., McMurray, M. S., Guzman, D. B., Nair, G., Shi, Y., McCormack, K. M., … Sanchez, M. M. (2017). Maternal buffering beyond glucocorticoids: impact of early life stress on corticolimbic circuits that control infant responses to novelty. Soc Neurosci, 12(1), 50–64. doi:10.1080/17470919.2016.1200481

Huang, T. N., Chuang, H. C., Chou, W. H., Chen, C. Y., Wang, H. F., Chou, S. J., & Hsueh, Y. P. (2014). Tbr1 haploinsufficiency impairs amygdalar axonal projections and results in cognitive abnormality. Nat Neurosci, 17(2), 240–247. doi:10.1038/nn.3626

Jimenez-Castellanos, J. (1949). The amygdaloid complex in monkey studies by reconstructional methods. J. Comp. Neurol., 91(3), 507–526, illust.

Kempermann, G., Jessberger, S., Steiner, B., & Kronenberg, G. (2004). Milestones of neuronal development in the adult hippocampus. Trends Neurosci., 27(8), 447–452.

Klüver, H., & Bucy, P. C. (1939). Preliminary analysis of functions of the temporal lobes in monkeys. Arch. Neurol. Psychiatry, 42, 979–997.

Kohler, S. J., Williams, N. I., Stanton, G. B., Cameron, J. L., & Greenough, W. T. (2011). Maturation time of new granule cells in the dentate gyrus of adult macaque monkeys exceeds six months. Proc. Natl. Acad. Sci. U. S. A., 108(25), 10326–10331. doi:1017099108 [pii] 10.1073/pnas.1017099108

LeDoux, J. E. (2000). Emotion circuits in the brain. Annu. Rev. Neurosci., 23, 155–184. doi:10.1146/annurev.neuro.23.1.155

McCormick, K., Gualano, M. F., Kerr, D., Rockcastle, N., & Cameron, J. L. (2005). Social bond disruption in early life has behavioral consequences which remain evident through puberty. Paper presented at the Society for Neuroscience, San Diego, CA.

Mihalas, A. B., & Hevner, R. F. (2017). Control of neuronal development by T-box genes in the brain. In Current Topics in Developmental Biology (Vol. 122): Elsevier Inc.

Nishijo, H., Ono, T., & Nishino, H. (1988). Topographic distribution of modality-specific amygdalar neurons in alert monkey. J. Neurosci., 8(10), 3556–3569.

O’Connor, T. G., & Rutter, M. (2000). Attachment disorder behavior following early severe deprivation: extension and longitudinal follow-up. English and Romanian Adoptees Study Team. J Am Acad Child Adolesc Psychiatry, 39(6), 703–712.

Purves, D., Snider, W. D., & Voyvodic, J. T. (1988). Trophic regulation of nerve cell morphology and innervation in the autonomic nervous system. Nature, 336(6195), 123–128. doi:10.1038/336123a0

Rosene, D. L., Roy, N. J., & Davis, B. J. (1986). A cryoprotection method that facilitates cutting frozen sections of whole monkey brains for histological and histochemical processing without freezing artifact. J. Histochem. Cytochem., Vol. 34, No. 10, 1301–1315.

Sabatini, M. J., Ebert, P., Lewis, D. A., Levitt, P., Cameron, J. L., & Mirnics, K. (2007). Amygdala gene expression correlates of social behavior in monkeys experiencing maternal separation. J. Neurosci., 27(12), 3295–3304.

Schumann, C. M., Bauman, M. D., & Amaral, D. G. (2011). Abnormal structure or function of the amygdala is a common component of neurodevelopmental disorders. Neuropsychologia, 49(4), 745–759. doi:10.1016/j.neuropsychologia.2010.09.028

Slomianka, L., & West, M. J. (2005). Estimators of the precision of stereological estimates: an example based on the CA1 pyramidal cell layer of rats. Neuroscience, 136(3), 757–767. doi:10.1016/j.neuroscience.2005.06.086

Sorrells, S. F., Paredes, M. F., Velmeshev, D., Herranz-Perez, V., Sandoval, K., Mayer, S., … Alvarez-Buylla, A. (2019). Immature excitatory neurons develop during adolescence in the human amygdala. Nat. Commun., 10(1), 2748. doi:10.1038/s41467-019-10765-1

Sullivan, R., Wilson, D. A., Feldon, J., Yee, B. K., Meyer, U., Richter-Levin, G., … Braun, K. (2006). The International Society for Developmental Psychobiology annual meeting symposium: Impact of early life experiences on brain and behavioral development. Dev. Psychobiol., 48(7), 583–602.

Suomi, S. J., Collins, M. L., & Harlow, H. F. (1973). Effects of permanent separation from mother on infant monkeys. Dev. Psychol., 9(3), 376–384. doi:10.1037/h0034896

Tottenham, N., Hare, T. A., Millner, A., Gilhooly, T., Zevin, J., & Casey, B. J. (2011). Elevated Amygdala Response to Faces Following Early Deprivation. Dev Sci, 14(2), 190–204. doi:10.1111/j.1467-7687.2010.00971.x

Tottenham, N., Hare, T. A., Quinn, B. T., McCarry, T. W., Nurse, M., Gilhooly, T., … Casey, B. J. (2010). Prolonged institutional rearing is associated with atypically large amygdala volume and difficulties in emotion regulation. Dev Sci, 13(1), 46–61. doi:DESC852 [pii] 10.1111/j.1467-7687.2009.00852.x

Ulfig, N., Setzer, M., & Bohl, J. (2003). Ontogeny of the human amygdala. Ann. N. Y. Acad. Sci., 985, 22–33.

West, M. J., Slomianka, L., & Gundersen, H. J. (1991). Unbiased stereological estimation of the total number of neurons in thesubdivisions of the rat hippocampus using the optical fractionator. Anat. Rec., 231(4), 482–497.

